# Towards understanding stomatal decoupling under extreme heat: integrating stomatal kinetics, leaf light responses, and the role of stomatal versus cuticular conductance

**DOI:** 10.64898/2026.10.24.760857

**Authors:** Renée M. Marchin, Alessandro Ossola, Mark G. Tjoelker, David S. Ellsworth

## Abstract

- Earth’s forests are increasingly exposed to hotter drought events, causing unprecedented dieback of trees and forests. Evidence of stomatal decoupling at high leaf temperatures (T_leaf_) is mounting, but the driving mechanisms and variability across species are unknown.
- We experimentally imposed heat and drought stress onto two rainforest and two semi-arid tree species in a glasshouse study to better understand accelerated water loss at high T_leaf_ by measuring plant thermal and hydraulic balance.
- Stomatal decoupling at high T_leaf_ occurred in two species, including the most anisohydric species, but did not occur in a vulnerable rainforest species. Accelerated leaf water loss occurred predominantly through open stomata, rather than the cuticle, but some species prioritised thermal acclimation over leaf thermoregulation during heat stress. This suggests decoupling occurs only in species with high capacity for transpirational cooling (>5°C), or alternatively, that full sunlight is required for stomatal decoupling.
- The maximum rate of light-induced stomatal opening was up to two times faster, while dark-induced stomatal closing was 14 minutes slower on average in leaves at 34°C, relative to 25°C. Stomatal kinetics are an overlooked trait that may help to better predict accelerated water loss under high T_leaf_ across species.

## Introduction

The last decade (2015–2025) has been the warmest in Earth’s climate record, with 2023, 2024, and 2025 being the three hottest years in recent history (WMO, 2026). Extreme temperatures >50 °C have been recorded in at least ten countries over the last several years, leading to increased interest in how plants cope with extreme heat stress (Kacprzyk *et al*., 2025). Trees have generally been thought to be resilient to acute heat stress, if sufficient water is available (Teskey *et al*., 2015), but recent extreme heat (>40 °C) events have caused crown dieback, reduced growth, and tree mortality in forests on multiple continents (Still *et al*., 2023; Gauthey *et al*., 2024; Xu *et al*., 2024). Most heatwaves tend to occur in areas where negative soil moisture anomalies are coupled to positive sensible heat flux anomalies (Bi *et al*., 2026). In recent decades, the dominant driver of these ‘compound heat and drought events’ or simply ‘hotter droughts’ has switched from rainfall deficit to excess heat (Alizadeh *et al*., 2020). There is growing recognition that the potential effects of heatwaves on forests have been underappreciated (Breshears *et al*., 2021). An improved understanding of the effects of heatwaves on tree physiology is needed, including the mechanisms driving leaf responses to heat in diverse tree species.

The dominant pathway for water loss from plant leaves is through stomata and is termed stomatal conductance (g_s_). Plants close their stomata to prevent excessive water loss under high evaporative demand (Lange *et al*., 1971; Mott & Parkhurst, 1991). The temperature effect on stomata is thus confounded by vapour pressure deficit (VPD), because warming causes an exponential increase in VPD (Grossiord *et al*., 2020). The direct temperature response of stomata can be determined in intact leaves under controlled conditions, though increasing temperature while maintaining constant VPD is difficult to achieve. The few careful studies that have done so found that, on average, g_s_ increases with increasing leaf temperature (T_leaf_) by about 2% per °C at constant VPD, relative to its value at 25 °C, equivalent to a doubling of g_s_ from 10 to 40 °C (Mills *et al*., 2024). The response varies considerably among species, with only half of species showing increased g_s_ with increasing T_leaf_ at constant VPD, and other species having negative, peaked, or little response of g_s_ to increasing T_leaf_ (Mills *et al*., 2024). At high temperatures >40 °C, a phenomenon known as ‘stomatal decoupling’ can occur, where net photosynthesis (A_net_) is inhibited and yet g_s_ continues at rates notably above zero (Drake *et al*., 2018; Aparecido *et al*., 2020; Marchin *et al*., 2023; Gauthey *et al*., 2024). A recent meta-analysis of 207 studies found that warming increased transpiration and decreased A_net_, while g_s_ exhibited no clear directional trend (Wang *et al*., 2026). Further research is needed to resolve our poor understanding of this decoupling response and improve predictions of carbon and water fluxes between the land and atmosphere (Blonder *et al*., 2023).

It is unclear what mechanisms control g_s_ and stomatal regulation in leaves at high temperatures (but see (Mills *et al*., 2024). Stomatal conductance has a net intrinsic temperature sensitivity due to physical changes in gas properties, according to Fick’s Law, Dalton’s Law, and the ideal gas law (Mills *et al*., 2024; Kowalski *et al*., 2025). At high T_leaf_, water vapour dynamics physically decouple leaf carbon and water fluxes – high humidity inside leaves boosts transpiration, while photosynthesis is inhibited by the simultaneous suppression of CO_2_ (Kowalski *et al*., 2025). Observed stomatal temperature responses are 4-12 times larger in magnitude than this purely physical response, indicating biological mechanisms are also involved (Mills *et al*., 2024). To actively mediate stomatal movements and aperture, the guard cell chloroplasts sense CO_2_ and perceive photosynthetically active radiation (Zeiger, 1983; Zeiger *et al*., 2002). Stomatal opening, or reopening, at high temperatures is promoted by blue light (Kostaki *et al*., 2020) and may function to support leaf cooling through latent heat dissipation (Urban *et al*., 2017; Marchin *et al*., 2022; Evans *et al*., 2025; Wang *et al*., 2026). Notably, the response time of stomata to a sudden increase in light (i.e., sunfleck) requires >20 minutes to reach maximal opening (Drake *et al*., 2013; Meinzer *et al*., 2017), which is an order of magnitude slower than the nearly instantaneous photosynthetic response (Kirschbaum *et al*., 1988; Lawson *et al*., 2018). No study has yet examined how warming may differentially affect stomatal kinetics across species.

Plants also lose water through leaky stomata and the waxy leaf cuticle, together termed minimum conductance (g_min_). Normally, g_min_ is a small fraction (mean: 4.89 ± 2.67 mmol^-2^ s^-1^; Duursma et al., 2019) of average g_s_ (∼250 mmol^-2^ s^-1^; (Murray *et al*., 2019), but g_min_ may approach or dominate tree water loss on hot days (Garen & Michaletz, 2025). The temperature response of g_min_ often exhibits a biphasic relationship, where sharp increases in g_min_ are observed above a breakpoint temperature, typically around 35-45 °C (Schreiber, 2001; Bueno *et al*., 2019; Billon *et al*., 2020; Slot *et al*., 2021). As with g_s_, the temperature effect on g_min_ is confounded by VPD, although most studies have not attempted to disentangle the two factors during measurement. One recent careful study increased temperature while maintaining constant VPD, finding that g_min_ of two tree species had a stronger temperature response, along with lower breakpoint temperatures, than under variable VPD (Garen *et al*., 2026). There is substantial variability in the g_min_ temperature response among species, with some species showing no temperature response (Slot *et al*., 2021) or declining g_min_ as temperature increases (Garen & Michaletz, 2025). Previous studies have been unable to resolve if stomatal decoupling is caused by accelerated water loss through open stomata (i.e., g_s_) or through closed stomata and the cuticle (i.e., g_min_).

A better understanding of the generality of stomatal decoupling across taxa and the driving factor for leaf water loss at high temperature (i.e., g_s_ versus g_min_) is needed to accurately model plant-atmosphere gas exchange. In this study, we aim to better understand accelerated water loss at high T_leaf_ by measuring plant water use, thermal regulation, and photosynthesis across four tree species exposed to concurrent experimental heat and drought stress in a glasshouse experiment. Half of the potted seedlings of two rainforest (*Acronychia oblongifolia*, *Pomaderris apetala*) and two semi-arid woodland tree species (*Brachychiton rupestris*, *Pittosporum angustifolium*) were exposed to a gradual, five-week experimental drought of moderate intensity (Marchin *et al*., 2020), before all plants were exposed to a six-day heatwave with a maximum air temperature of 42°C (Supporting Information Fig. S1). In a previous study, we identified evidence of stomatal decoupling for some study species during the experimental heatwave (Marchin *et al*., 2023). To better understand the mechanisms and consequences of this differential response across species, we addressed the following questions:

1. How are plant water use and leaf thermoregulation under drought stress related to vulnerability to heatwaves across species?
2. What is the driving pathway for leaf water loss at high T_leaf_ – g_s_ or g_min_?
3. How does warming affect light-induced stomatal kinetics across tree species from diverging biomes?

## Materials and Methods

### Study species and glasshouse experiment

Four native Australian tree species were selected for study; two mesic species originating from rainforests, and two xeric species originating from semi-arid woodlands (Table 1; Supplementary Table S1). Planting stock (*n*=10 plants per species) ranged from tubestock to 140-mm pot size and was obtained from commercial nurseries in Australia located near the centroid of the species’ native range. Seedlings were bare-rooted and transplanted into 6-L square pots containing native potting mix (<30% sand/coir, >70% screened composted pine bark; Australian Growing Solutions, Tyabb, VIC, Australia), 38 g of controlled-release native plant fertiliser (Scotts Australia Osmocote Slow Release, Bella Vista, NSW, Australia), and 1.25 g of systemic insecticide and fertiliser tablet (Yates Confidor, Padstow, NSW, Australia).

**Table 1.** Four native Australian tree species selected for this study, including their growth habit, natural occurrence, leaf size, leaf mass per area, and wood density. Values are means of 16-20 leaves or 5 stems per species; means not connected by the same letter are significantly different (Tukey honestly significant difference, *p*<0.05). Xeric species are shaded.

| Species | Growth habit | Natural occurrence | Leaf size (cm<sup>2</sup>) |  |  | LMA (g m<sup>-2</sup>) |  |  | Wood density (g cm<sup>-3</sup>) |  |  |
| --- | --- | --- | --- | --- | --- | --- | --- | --- | --- | --- | --- |
| *Acronychia oblongifolia* | Evergreen | warm coastal rainforest | 21.9 | ± 1.7 | c | 75.2 | ± 1.9 | b | 0.487 | ± 0.025 | a |
| *Pomaderris apetala* | Evergreen | wet sclerophyll forest | 24.2 | ± 3.7 | c | 57.3 | ± 3.3 | a | 0.438 | ± 0.028 | ab |
| *Brachychiton rupestris* | Drought deciduous | semi-arid bottle tree scrub | 8.5 | ± 0.6 | a | 77.1 | ± 3.1 | b | 0.368 | ± 0.024 | b |
| *Pittosporum angustifolium* | Evergreen | arid mallee woodland | 13.6 | ± 1.5 | b | 126.6 | ± 6.0 | c | 0.494 | ± 0.020 | a |

Plants were grown in a controlled environment glasshouse at the Hawkesbury Institute for the Environment (Western Sydney University, Richmond, NSW) from 1 October 2018 to 1 February 2019. Seedlings were randomly rotated within and between glasshouse bays monthly to allow uniform solar irradiance for growth. Seedlings were well-watered using drip irrigation for 6 or 11 weeks to establish roots, allow formation of new leaves, and acclimate to the glasshouse environment. Average glasshouse temperature was 28 °C with a diurnal range from 22-35 °C (Supporting Information, Fig. S1) to represent summer conditions in southeastern Australia. Daily maximum photosynthetically active radiation (PAR) was >2000 µmol m^-2^ s^-1^, daytime relative humidity was 40-95%, and daytime VPD was 0.2-3 kPa inside the glasshouse (Supporting Information, Fig. S1). Watering to saturation required 1 L at 6:00 at the beginning of the experiment and was increased to a maximum of 4.5 L daily (delivered at 8:00, 13:00, and 17:00) as plants grew larger.

After the acclimation period, half of the plants (*n*=5 plants per species) were exposed to a gradual, five-week drought (Supporting Information, Fig. S2) following the method described in Marchin et al. (2020). The remaining half served as well-watered controls. Briefly, pots were placed inside 100-L plastic containers on top of a 23-cm column of commercial porous foam (Oasis IDEAL Floral Foam Maxlife brick; Smithers-Oasis) with an adjustable water table. Depth to the water table was progressively increased from 1 cm (on day 1) to 15 cm (on day 8) to 22 cm (on day 15), where it was maintained for an additional 3 weeks (until day 35). Water levels were checked daily and maintained within ±1-cm of the target level. The target drought intensity was a volumetric water content (VWC) of 7.5 ± 2.5%, which was well below the permanent wilting point of the soil (14%). The soil VWC was monitored in the morning (8:00-10:00) using a 20-cm soil water content probe (CS658 HydroSenseII, Campbell Scientific Inc., Logan, UT, USA); soil VWC was measured weekly for the first three weeks of experimental drought, but every three days thereafter during plant measurement weeks. Further details can be found in Marchin et al. (2020).

In the final week of water deficit (days 29-35), all plants (droughted and irrigated) were exposed to a six-day heatwave by moving pots into another glasshouse bay. The average heatwave temperature was 35 °C (i.e., +7 °C above baseline temperature), with a diurnal range of 31-42 °C and the maximum temperature spanning 2 h at midday (12:00-14:00; Supporting Information, Fig. S1). The daytime (10:00-16:00) relative humidity during the heatwave was 57%, which was +4% above baseline relative humidity (61%; Fig. S1C). During the heatwave, control pots (*n*=5 plants per species) were well-watered using drip irrigation (mean VWC: 32%, ranging from 23-36%), while drought pots were maintained on foam with 22-cm depth to the water table (VWC=3-9%; Supporting Information, Fig. S2). It was not possible to measure all species in the same week, so *B. rupestris* and *P. apetala* were exposed to the heatwave in the austral summer from 7-12 December 2018, while *A. oblongifolia* and *P. angustifolium* were exposed to the heatwave from 11-16 January 2019. Only three of six heatwave days were sunny in December 2018, while five of six heatwave days were sunny in January 2019 (Supporting Information, Fig. S3).

### Plant functional traits

We measured plant functional traits related to leaf temperature regulation, including leaf size, leaf mass per area (LMA), and the cooling capacity of transpiration, plus wood density. Leaf size was measured for two fully-expanded leaves per plant (*n*=9-10 plants per species) using a flatbed scanner and ImageJ software (Schneider *et al*., 2012). Leaves were then oven-dried for 48 hours at 70 °C to obtain dry mass. To measure the capacity for transpirational cooling, we applied an even layer of Vaseline to the abaxial side of two leaves per plant (or leaflets, for the compound leaves of *B. rupestris*) as antitranspirant. All species are hypostomatous, so this has been shown to reduce transpiration by 95-98% (Curtis, 1936). We then measured the temperature of the Vaseline leaf and the opposite transpiring leaf, or neighbouring leaflet, using an infrared camera (FLIR T640, Wilsonville, OR, USA). Measurements occurred at midday (12:00-16:00) on sunny days in January and February 2019, after the recovery period, while plants were well-watered and measured leaves were exposed to full sun. Wood density was determined at the end of the experiment for five plants per species by splitting a 5-cm stem segment to remove the pith and bark. Fresh sapwood volume was determined using the water displacement method. Wood samples were then oven-dried to constant mass at 105 °C.

### Leaf gas exchange and plant water relations

Leaf-level gas exchange was measured on two fully-expanded leaves per plant using a portable infrared gas analyser equipped with a red–blue light source (LI-6400XT, LI-COR Biosciences, Lincoln, NE, USA). Leaves were measured around the hottest time of day, from 10:00 to 13:30 as per experimental settings. Measurements were collected under average summer air temperatures (27-35 °C) at the beginning, middle, and end of the drought, plus on two recovery days after the heatwave (Supporting Information, Fig. S4). Measurements were also collected under heatwave air temperatures (36-42 °C) on days 1, 3, and 5 of the heatwave (Supporting Information, Fig. S4).

Net photosynthesis (A_net_) and stomatal conductance to water vapour were recorded for a period of 30 s after conditions inside the cuvette stabilized (usually 1-2 min) and then averaged for each leaf. Here, we have inferred the measured water vapour fluxes are via the stomata and use the term *g*_s_, but leaf transpiration is a combination of water lost through the stomata and desorbed from the leaf surface through the cuticle. Measurements were made under saturating light (PAR=1800 μmol m^−2^ s^−1^), ambient leaf temperature, CO_2_ concentration of 420 ppm, and relative humidity ±10% of ambient. Leaf area was measured using a flatbed scanner and ImageJ software (https://imagej.nih.gov/ij/, (Schneider *et al*., 2012) for area corrections to the gas exchange measurements for xeric species with thin leaves that did not fill the cuvette.

The same leaves were used to determine the midday leaf water potential (Ψ_mid_) with a pressure chamber (Model 1505D; PMS Instruments, Albury, OR, USA) by measuring two leaves per plant (*n*=10 plants per species) at the peak of drought (baseline, drought day 26) and also during the heatwave (drought day 33). Leaves were stored inside a sealed, humidified plastic bag containing a small piece of wet paper towel and kept cool and dark until equilibrated for approximately 1 h, with measurements completed within 3 h of sample collection. A dissecting microscope was placed above the pressure chamber, with fiber-optic lights, to clearly observe the balance pressure, which was recorded as the mid-point between the first sign of water from the petiole and when the entire xylem area is wetted.

To compare levels of drought stress across species, Ψ_mid_ was compared to the leaf water potential at turgor loss point (π_tlp_) for each species. The osmotic potential of fully-expanded, fully-hydrated leaves (*π*_o_; *n*=9-10 plants per species) was measured using an osmometer (WP4C Dewpoint PotentiaMeter; Decagon Devices), according to the method described by Bartlett et al. (2012). Briefly, leaves were collected and rehydrated overnight for 12 h using the standing rehydration method (Arndt *et al*., 2015) to ensure fully-hydrated leaves (i.e. Ψ_leaf_ ≥‒ 0.3 MPa) were used for comparison. The midrib was removed, leaves were frozen in liquid nitrogen, equilibrated for 10 min, and punctured with sharp-tipped forceps before measurement. Measurements were recorded for 20-30 min until equilibrium was reached as indicated by <0.01 MPa change over 2 min. Osmometer measurements of *π*_o_ were used to estimate species’ π_tlp_ using the equation from Bartlett et al. (2012): *π_tlp_* = 0.832*π_o_* − 0.631.

### Minimum conductance

The temperature response of minimum conductance (g_min_) was measured following the ‘bench drying’ method described by Sack *et al*. (2003). At the end of the experiment, fully-developed leaves were harvested and rehydrated overnight for 12 h before wrapping the cut petiole with parafilm to minimize water loss through the petiole. Leaves were placed inside a ventilated climate chamber with 330 μmol m^-2^ s^-1^ PAR and slowly desiccated at five air temperatures: 20, 27, 34, 40, and 45 °C. Air temperature was monitored with the built-in growth chamber sensor; relative humidity was measured with a HOBO MX1101 sensor and ranged from 67% at 20 °C to 59% at 45 °C. Leaves were allowed to dehydrate until stomata were closed (0.5-4 h) and then weighed every 15-25 minutes, depending on the temperature, for a period of 3 hours. Leaves were stored in plastic bags to prevent water loss when not being dried or weighed. Leaf area was measured at the beginning and end of the dehydration process using a flatbed scanner and ImageJ software (https://imagej.nih.gov/ij/, (Schneider *et al*., 2012) to account for leaf shrinkage during desiccation. The g_min_ (*n*=5-15 leaves per temperature) was calculated in units of mol m^-2^ s^-1^ for comparison with g_s_, after removal of the initial nonlinear portion of the mass loss curve (i.e., when stomata were not fully closed and contributing to water loss), retaining linear relationships containing 4-8 mass measurements for each leaf. The g_min_ was calculated as rate of water loss per unit leaf area per unit time, divided by VPD, calculated from air temperature, relative humidity, and atmospheric pressure.

### Leaf temperature and leaf critical temperature

Leaf temperature was measured for one leaf per plant using fine-wire thermocouples (36-gauge Type T; Omega) attached to the abaxial leaf surface with surgical tape; measurements were recorded every 1 min by an automated datalogger (CR1000; Campbell Scientific Inc.). Leaf critical temperature (T_crit_) was measured for control and droughted plants (*n*=3-5 plants per treatment) under baseline and heatwave conditions. The temperature-dependent rise of steady-state chlorophyll *a* fluorescence (T_leaf_ − F_o_) was measured *in vivo* on dark-adapted leaves following the method of Schreiber and Berry (1977), with several modifications described in (Cox *et al*., 2026). First, plants were temporarily relocated into the laboratory, and fully-expanded leaves were dark-adapted for at least 10 min or until a stable fluorescence signal was achieved. Second, attached leaves were flattened and pressed firmly onto a filter paper on top of a Peltier thermoelectric cooler (Model APH-161-12-18-E; European Thermodynamics) inside a custom-built leaf cuvette. Leaves were progressively warmed, and the F_o_ was recorded every 1 s by a fluorometer (MINI-PAM-II/B; Heinz Walz GmbH, Effeltrich, Germany) as Peltier temperature in the cuvette increased from 35 to 70 °C at a rate of 1 °C min^−1^. The T_crit_ was calculated as the intersection of linear slow- and fast-rise phases of T_leaf_ − F_o_ curves, where minimum F_o_ was averaged from 35 to 38 °C using a zero-slope line for the slow-rise phase, and the fast-rise phase was calculated using F_o_ from ±1.5 min of the midpoint between minimum and maximum F_o_ values.

### Plant thermal damage

We measured two damage indicators to assess heat tolerance: the maximum quantum yield of PSII (F_v_/F_m_) and whole-plant crown dieback. To assess F_v_/F_m_, variable (F_v_) and maximum chlorophyll *a* fluorescence (F_m_) were measured with a fluorometer (MINI-PAM-II/B; Heinz Walz GmbH) on one whole dark-adapted leaf per plant using dark adaptation leaf clips (DLC-8; Heinz Walz GmbH). Leaves (*n*=4-5 plants per species) were selected and flagged to permit repeat comparison under baseline and heatwave conditions. Prior to F_v_/F_m_ measurement, the leaves were dark adapted for a minimum of 30 min. After measuring F_o_, a saturation pulse was applied to measure maximum fluorescence (F_m_). The difference in F_v_/F_m_ between baseline and heatwave treatments (ΔF_v_/F_m_) was calculated to estimate the level of heat-induced chronic photoinhibition; baseline measurements were made the day before the heatwave, while heatwave measurements were made on the fifth day of the heatwave.

Whole-plant crown dieback was assessed visually by estimating the percentage of leaf scorch or necrosis of leaves, relative to total leaf area. Two expert observers provided independent estimates under baseline conditions (*D*_1_, *D*_2_), after the heatwave (*D*_HW1_, *D*_HW2_), and after two weeks of recovery (*D*_Rec1_, *D*_Rec2_). Crown dieback was calculated as (*D*_max1_ + *D*_max2_/2) – (*D*_1_ + *D*_2_/2) for each plant, where data from both observers was averaged and maximum dieback (*D*_max1_, *D*_max2_; either after the heatwave or after rewatering) was used to account for delayed heatwave effects.

### Stomatal kinetics

To determine the temperature effect on the rate of light-induced stomatal opening, we followed a protocol similar to that described by Drake *et al*. (2013). We tracked time courses of light-induced stomatal opening and subsequent stomatal closing, which mimics stomatal responses during daytime sunflecks, at two different leaf temperatures: 25 and 34 °C. Plants were measured in the glasshouse at 34 °C or were relocated to the laboratory for measurement at 25 °C, after the end of the experiment. Measurements were conducted on mature leaves (*n*=4-8 leaves per species) from well-watered plants using a portable infrared gas analyser equipped with a red–blue light source (LI-6400XT, LI-COR Biosciences). Leaves were measured under ambient CO_2_ (420 ppm) and VPD of 1–2.2 kPa for 25 °C or 1.4–3.2 kPa for 34 °C. For light-induced stomatal opening curves, leaves were measured every 30 s for 20-30 min until stabilisation under low light (PAR=50 μmol m^−2^ s^−1^), representing shade conditions, before switching to saturating light (PAR=1800 μmol m^−2^ s^−1^) and continuing to measure every 2 min until g_s_ was stable for three consecutive measurements.

The same leaves were then measured to obtain stomatal closing curves by switching from saturating to low light (PAR=50 μmol m^−2^ s^−1^), with measurement every 2 min until g_s_ was stable for at least three consecutive measurements. For xeric species with thin leaves that did not fill the cuvette, leaf area corrections were applied to gas exchange measurements. Stomatal kinetics parameters included: g_s_ in the shade (g_low_), steady-state operating g_s_ in saturating light (g_op_), light-saturated photosynthesis (A_sat_), time taken to reach 50% of the point between g_low_ and g_op_ (t_50_), and the maximum rate of change in g_s_ during stomatal opening/closing ([Δg_s_/Δt]_max_).

### Statistical analyses

Species differences in plant functional traits, including leaf size, LMA, wood density, A_sat_, transpirational cooling, and π_tlp_ were analysed using one-way analyses of variance (ANOVA) with species as the fixed effect. Differences among species were analysed using Tukey’s Honestly Significant Difference (HSD) test.

To test whether the drought treatment affected *g*_s_, a repeated measures mixed model ANOVA was used (Bates *et al*., 2015); species and treatment were included as fixed effects and plant identity was nested within date as a random effect. The combined effect of the heatwave and drought on Ψ_mid_, T_crit_, ΔF_v_/F_m_, and maximum crown dieback was determined using full-factorial, mixed-model ANOVAs with species and treatment as the main effects; species identity was included as a random effect and treatment as a fixed effect. Separate models were used to test for species × treatment interactions, and if significant (*p*≤0.05), differences among species were analysed using Tukey’s HSD test. The relationship between leaf and air temperature during the heatwave was determined for a representative droughted plant for each species. Linear and quadratic regression models were tested, and the best model was selected using Akaike’s Information Criterion (AIC).

To determine if stomatal decoupling had occurred, we compared the change in A_net_ and g_s_ between ‘baseline’ (drought day 26) and ‘heatwave’ (drought days 28, 31, 33) conditions. These four measurement dates were used to ensure all plants were under peak drought treatment (Supporting Information Fig. S4). Stomatal decoupling was considered to occur if g_s_ did not decrease at high air temperatures (>40°C) in coordination with declines in A_net_ (Supporting Information Fig. S5). The temperature effect was quantified separately for A_net_ and g_s_ for control and drought treatments using mixed model ANOVAs with temperature (i.e., baseline vs. heatwave) as the main fixed effect; plant identity was included as a random effect. Thus, decoupling was considered to occur if, on hot days, g_s_ significantly increased (or did not change) while A_net_ significantly decreased.

We also used a more precise test based on the unified stomatal optimisation (USO) model to detect decoupling, as described by Medlyn et al. (2011):

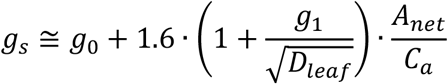

where D_leaf_ is leaf-to-air vapour pressure deficit, A_net_ is net photosynthesis, and C_a_ is atmospheric CO_2_ concentration. The model can be used to predict the expected change in g_s_ with A_net_ under heatwave conditions. The model includes two constants, g_0_ (the g_s_ at zero photosynthesis) and g_1_ (the slope parameter, inversely related to plant water use efficiency), which can be fitted to the model using the PLANTECOPHYS package in R (Duursma, 2015). However, the constant g_0_ can only be estimated accurately when photosynthesis approaches zero in the dataset, otherwise spurious values may be obtained (Duursma *et al*., 2019). We did not have sufficient measurements at zero photosynthesis to estimate g_0_ and hence fitted g_1_ only; g_1_ was fitted for control and droughted plants using all gas exchange data collected under average summer air temperatures (27–35 °C). We then predicted values under heatwave conditions for comparison to observations; linear regressions with zero intercepts were used to test whether the observed data departed significantly from the model. We tested both linear or quadratic regression models and selected the best model using AIC, corrected for small sizes.

The temperature response of g_min_ was analyzed for individual species using linear, quadratic, or piecewise regression analyses using the *segmented* package in R (Muggeo, 2008), after confirming that there was a T_air_ × species interaction. The best model (linear, quadratic, or bi-phasic) was selected using AIC, corrected for small sizes. The temperature response of stomatal kinetics parameters g_low_, g_op_, [Δg_s_/Δt]_max_, and t_50_ was analysed separately for opening and closing curves using mixed model ANOVAs with species and temperature (i.e., 25 vs. 34 °C) as the main fixed effects; plant identity was included as a random effect. To further compare the temperature effect on stomatal kinetics parameters [Δg_s_/Δt]_max_ and t_50_, mixed model ANOVAs with temperature and type (i.e., opening vs. closing) as the main fixed effects were used; plant identity was included as a random effect. Pairwise comparisons and significant interaction effects were explored using the function ‘emmeans’ (Lenth *et al*., 2018).

## Results

### Species differences in plant functional traits and drought strategy

The two rainforest species *A. oblongifolia* and *P. apetala* had larger leaves (*F*_3,35_=17.86, *p*<0.001) with lower LMA (*F*_3,35_=64.93, *p*<0.001) and lower A_sat_ (*F*_3,49_=17.61, *p*<0.001), relative to the two xeric species *B. rupestris* and *P. angustifolium* (Tables 1, 2). The highest π_tlp_ was found for *A. oblongifolia*, while *P. angustifolium* had the lowest π_tlp_ (*F*_3,34_=27.89, *p*<0.001; Table 2). Wood density was similar for all evergreen species, while the drought-deciduous bottle tree (*B. rupestris*) had the lowest wood density (*F*_3,17_=5.24, *p*=0.010; Table 1). Two species, *A. oblongifolia* and *P. angustifolium*, had a greater capacity for transpirational cooling at 5-6 °C under summer temperatures (T_air_= 35 °C), relative to the other two species, *P. apetala* and *B. rupestris* (∼2 °C; *F*_3,73_=20.38, *p*<0.001; Table 2).

**Table 2.**
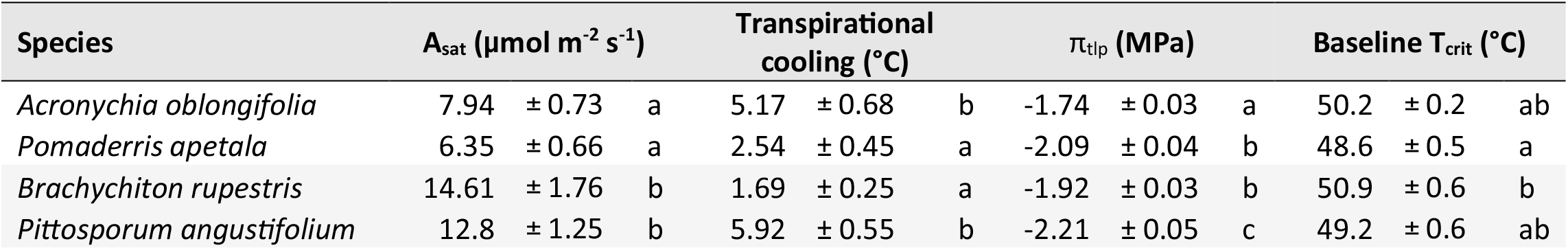
Species differences in light-saturated net photosynthesis (A_sat_), the capacity for transpirational cooling under summer temperatures (T_air_= 35 °C), leaf water potential at turgor loss point (π_tlp_), and baseline leaf critical temperature (T_crit_). Values are means of 11-20 leaves per species; means not connected by the same letter are significantly different (Tukey honestly significant difference, *p*<0.05). Xeric species are shaded.

Soil VWC reached our target drought level around drought day 15 (Supporting Information Fig. S2), resulting in a moderate drought treatment that significantly reduced g_s_ of all species (*F*_1,315_=121.25, *p*<0.001; Supporting Information Fig. S4). Due to species’ different drought tolerances and their dynamic regulation of g_s_ and Ψ_mid_ (*F*_3,32_=8.36, *p*<0.001), drought significantly decreased Ψ_mid_ (*F*_1,32_=9.68, *p*=0.004) of the two rainforest species but not the most isohydric species (*B. rupestris*; Fig. 1A). All species except the most anisohydric species (*P. angustifolium*) maintained their Ψ_mid_ above their mean π_tlp_, at least under average summer temperatures (Fig. 1A). The combined effect of the heatwave and drought on Ψ_mid_ differed among species (*F*_3,32_=26.39, *p*<0.001; Fig. 1C). For all species except the isohydric *B. rupestris*, Ψ_mid_ of droughted plants fell below their mean π_tlp_ during the heatwave (Fig. 1C). For the anisohydric *P. angustifolium*, the Ψ_mid_ of all plants, including well-watered controls, exceeded their π_tlp_ during the heatwave (Fig. 1C).

**Figure 1.**
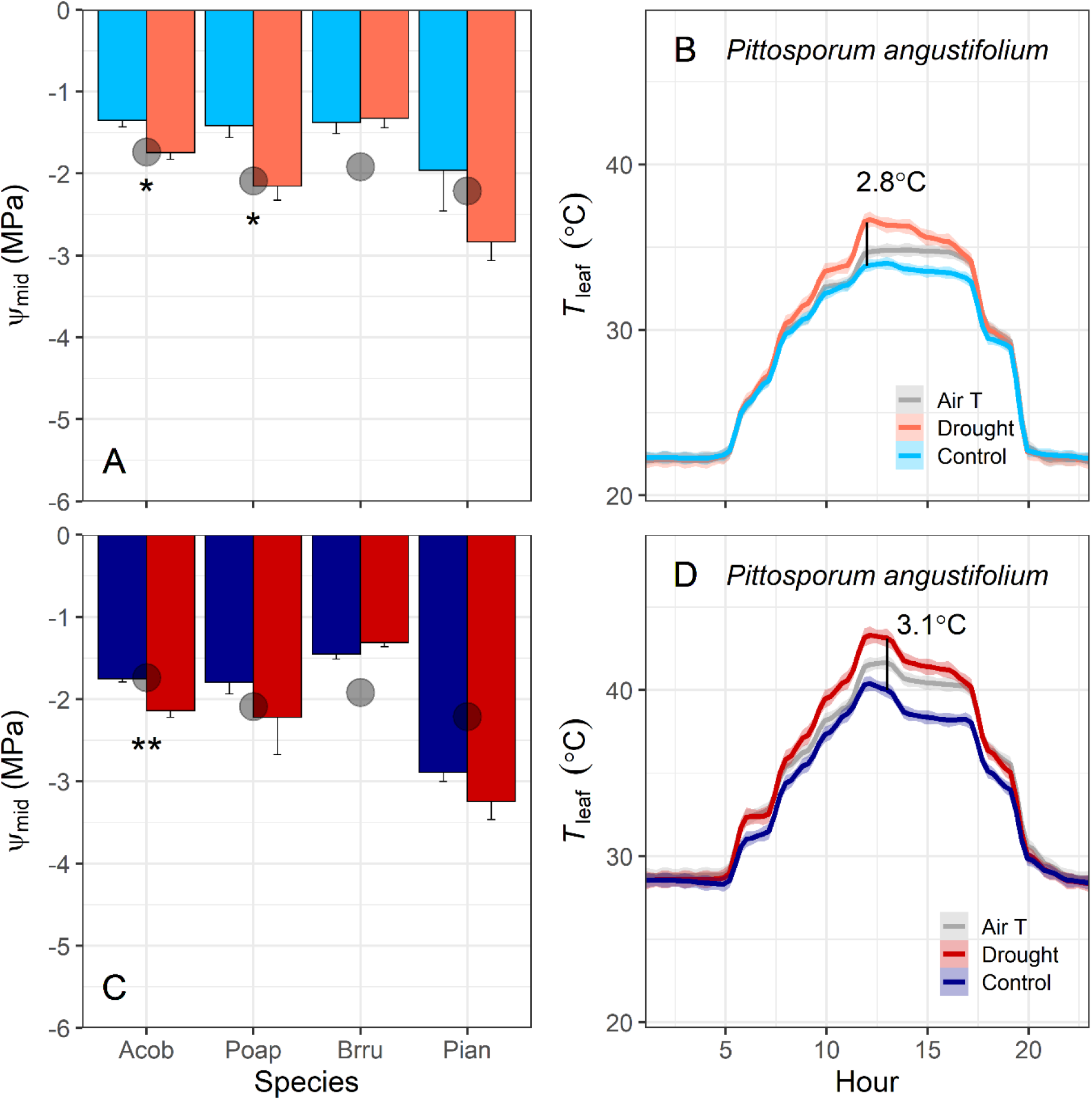
The effect of drought alone on (A) midday leaf water potential (Ψ_mid_, MPa) of four tree species (Acob: *Acronychia oblongifolia*, Poap: *Pomaderris apetala*, Brru: *Brachychiton rupestris*, Pian: *Pittosporum angustifolium*) and (B) diurnal patterns of leaf temperature (T_leaf_) in *P. angustifolium*. In panels C and D, the combined effects of drought plus heatwave are shown. The gray points in panels A and C indicate species mean leaf water potential at turgor loss point (π_tlp_). Values are means of 4-5 plants per treatment, and error bars indicate SE (unidirectional SE are presented for clarity). Asterisks denote significant differences between treatments: * *p*<0.05; ** *p*<0.01; *** *p*<0.001. In panels B and D, air temperature is plotted for reference (gray line), and the black vertical line indicates the mean temperature difference at midday between 5 plants per treatment; shaded lines indicate SE.

### Plant thermal regulation and tolerance under heat and drought stress

The T_leaf_ of droughted trees typically surpassed air temperatures during peak heatwave hours (10:00 to 16:00), while the T_leaf_ of well-watered control trees remained consistently below air temperature (Fig. 1B, 1D). For *P. angustifolium*, control leaves were an average of about 3 °C cooler at midday than droughted leaves (Fig. 1B, 1D), and droughted leaves in full sun reached >5 °C above air temperature (Fig. 2B). The largest differences between leaf and air temperatures were observed in a droughted *P. apetala* plant, whose hottest recorded T_leaf_ was >12 °C above T_air_ (Fig. 2C). The relationships between leaf and air temperature did not follow the 1:1 line, with droughted leaves of all species becoming hotter than air temperature above 40 °C (Fig. 2). This was most pronounced for the rainforest species *P. apetala*, whose leaf-to-air relationship was strongly nonlinear (*r*^2^=0.59, *p*<0.001; Fig. 2C).

**Figure 2.**
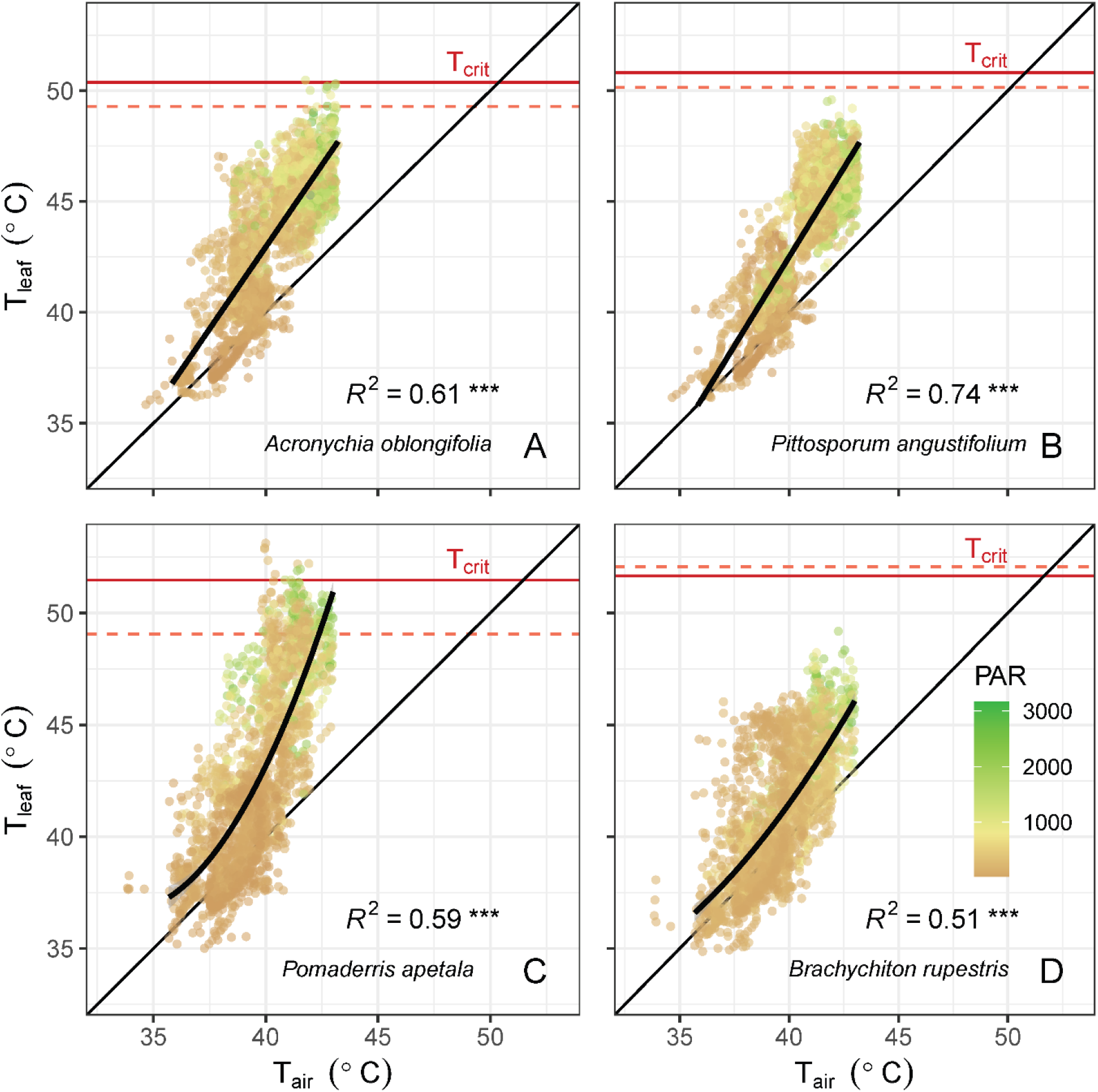
Relationship between daytime leaf temperature and air temperature for a droughted plant of (A) *Acronychia oblongifolia*, (B) *Pittosporum angustifolium*, (C) *Pomaderris apetala*, and (D) *Brachychiton rupestris*. Points were recorded every minute from 10:00 to 16:00 throughout the heatwave and are shaded by the photosynthetically active radiation (PAR, mol m^-2^ s^-1^) at time of measurement. Solid lines show linear or nonlinear regressions fit to the measured data. The 1:1 line is shown for reference. Asterisks denote significant relationships: ** *p*<0.01; *** *p*<0.001. Leaf critical temperature (T_crit_; horizontal lines) is shown for each plant; measurements were made two days before the heatwave (dashed line) and on the fifth day of the heatwave (solid line).

The T_crit_ differed among species (*F*_3,32_=5.35, *p*=0.004) and by treatment (*F*_1,32_=6.20, *p*=0.018), with droughted plants having significantly higher T_crit_ than control plants (mean of 50.5 ±0.3 °C vs. 48.8 ±0.4 °C, respectively). Among species, mesic *P. apetala* had lower T_crit_ than xeric *B. rupestris* under baseline conditions (Table 2). The T_crit_ of all plants acclimated to the heatwave, with control plants achieving similar critical thresholds as droughted plants (mean: 50.6 ±0.4 °C vs. 51.3 ±0.2 °C, respectively) with no difference among species (means: 50.1 ±0.5 °C to 51.5 ±0.5). The hottest leaves of both rainforest species approached or surpassed their T_crit_ under heat and drought stress, while T_leaf_ of the two xeric species remained below their critical thresholds (Fig. 2).

There was no mortality during the experiment but moderate levels of crown dieback (>20%) for three *P. apetala* tree seedlings (Fig. 3A). The effect of the heatwave on crown dieback differed among species (*F*_3,32_=5.35, *p*=0.004) and by treatment (*F*_1,32_=6.20, *p*=0.018), with droughted plants having significantly higher crown dieback than control plants (mean: 7.9 ±2.4% vs. 4.2 ±2.7%, respectively). Crown dieback was significantly higher in the rainforest species *P. apetala* than in the two xeric species, *P. angustifolium* and *B. rupestris* (Fig. 3A). The effect of the heatwave on ΔF_v_/F_m_ differed by species (*F*_3,32_=13.23, *p*<0.001) but not between control and droughted plants (*F*_1,32_=0.47, *p*=0.497). The heatwave decreased mean F_v_/F_m_ for the two species that underwent their experimental heatwave in a sunnier week, *A. oblongifolia* and *P. angustifolium*, but had no effect on mean F_v_/F_m_ for the other two species (Fig. 3B).

**Figure 3.**
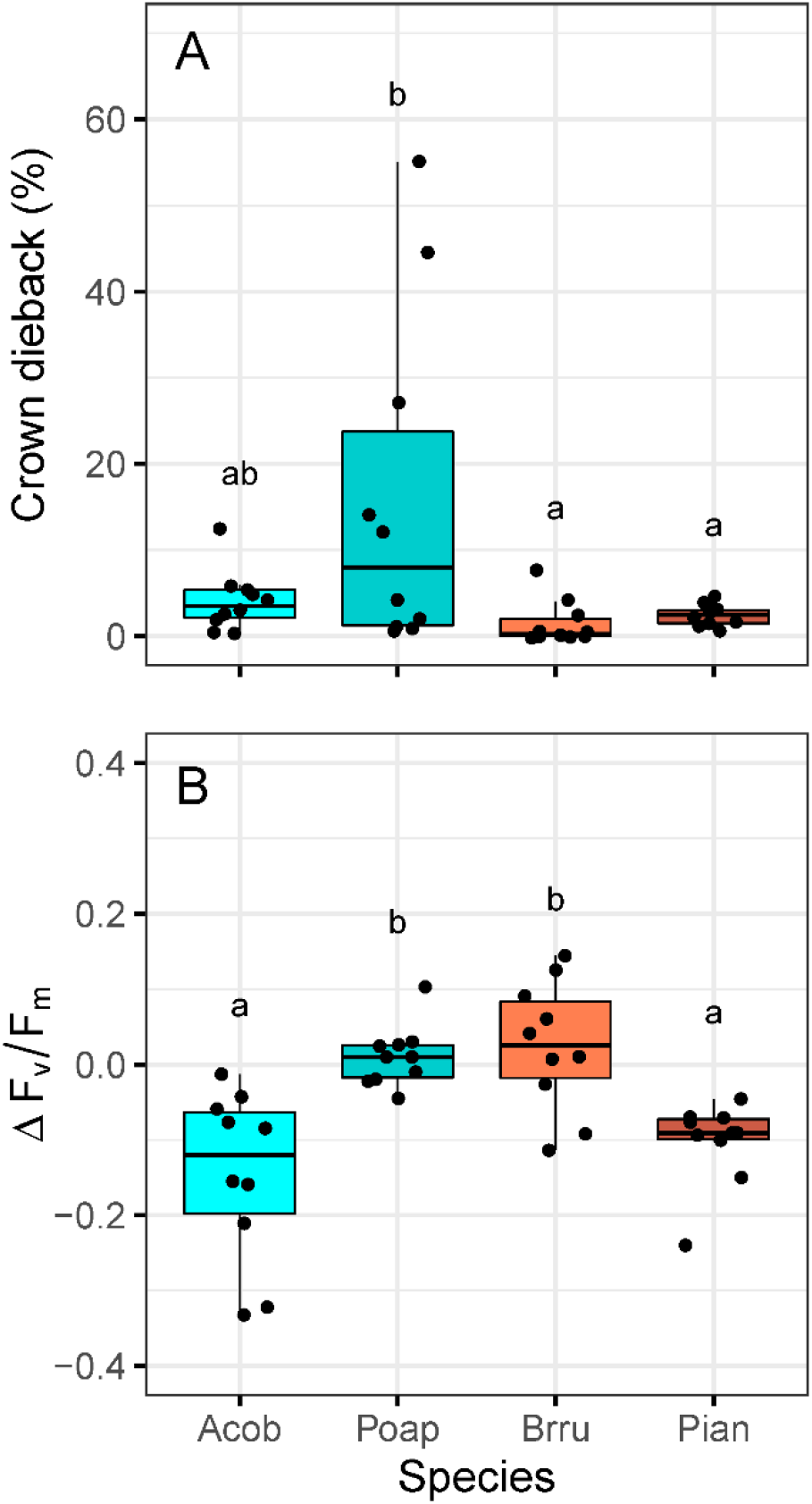
Species differences in (A) maximum crown dieback and (B) the difference between maximum quantum yield of PSII (Δ*F_v_/F_m_*) after the heatwave for control and droughted plants of four species (Acob: *Acronychia oblongifolia*, Pian: *Pittosporum angustifolium*, Poap: *Pomaderris apetala*, Brru: *Brachychiton rupestris*). Negative Δ*F_v_/F_m_* indicates heat-induced chronic photoinhibition on the fifth day of the heatwave. Points are individual plants (*n*=10 plants per species). Means not connected by the same letter are significantly different (Tukey honestly significant difference, *p*<0.05).

### Temperature effect on stomatal conductance to identify stomatal decoupling

The xeric species *P. angustifolium* had the strongest stomatal decoupling response to heat, which was evident in both control (Fig. 4B) and droughted plants (Supporting Information Fig. S6B), although mean A_net_ remained above zero (Supporting Information Fig. S5). Evidence for stomatal decoupling was also found for control plants of the rainforest species *A. oblongifolia* (Fig. 4A). Control plants significantly increased g_s_ during the heatwave (*P. angustifolium*: *F*_1,40_=4.15, *p*=0.049; *A. oblongifolia*: *F*_1,48_=4.26, *p*=0.045), despite a decrease in A_net_ at high air temperature (*P. angustifolium*: *F*_1,40_=5.78, *p*=0.022; *A. oblongifolia*: *F*_1,49_=13.66, *p*<0.001; Supporting Information Fig. S5). In addition, the observed g_s_ during heatwave conditions differed strongly from that predicted by the USO fitted to control plants under average summer temperatures (Fig. 4A, 4B) – the slope of the relationship between observed and predicted g_s_ was 1.40 for *P. angustifolium* and 1.62 for *A. oblongifolia* (Table 3).

**Figure 4.**
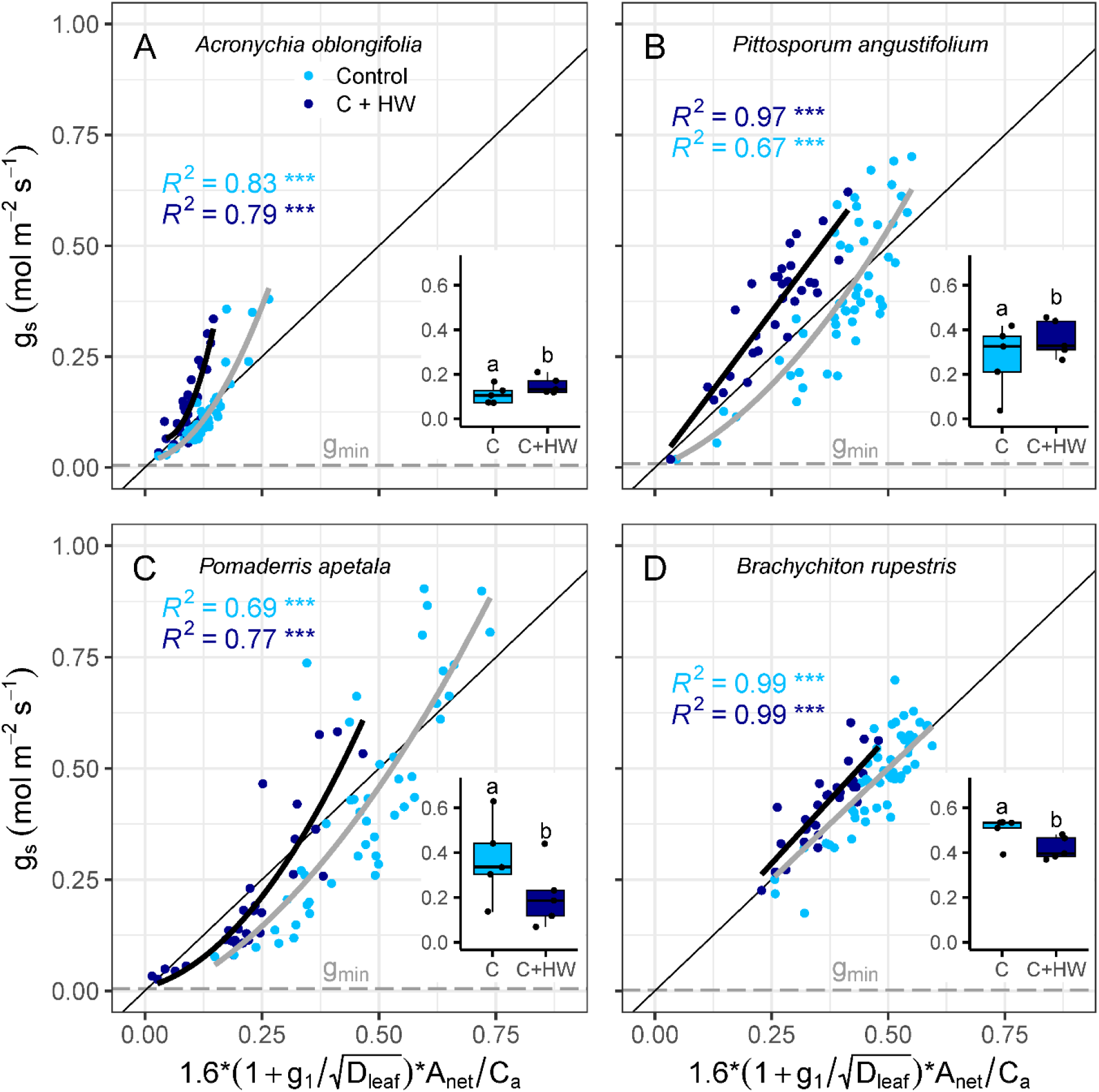
Relationships between observed stomatal conductance (g_s_) and the parameterised unified stomatal optimisation model (USO) for control plants of (A) *Acronychia oblongifolia*, (B) *Pittosporum angustifolium*, (C) *Pomaderris apetala*, and (D) *Brachychiton rupestris*. Responses are shown before (light blue) and during the heatwave (dark blue). Solid lines show linear or nonlinear regressions fit to the measured data. The 1:1 line between predicted and observed data is shown for reference. Points are 30-sec means for each leaf. Asterisks denote significant relationships: ** *p*<0.01; *** *p*<0.001. Insets show heatwave effects, where points are plant means. Means not connected by the same letter are significantly different (Tukey honestly significant difference, *p*<0.05). Maximum mean minimum conductance (g_min_; horizontal dashed line) is shown for each species (i.e., at 45 °C for panels A and B, at 27 °C for panels C and D).

**Table 3.** Summary of tests for whether the observed g_s_ data departed significantly from the unified stomatal optimization (USO) model, fitted separately for control and droughted plants using gas exchange data collected under average summer air temperatures (27–35 °C). Slopes (and 95% confidence intervals) are provided for the linear regressions between observed and predicted g_s_; significant deviations from the model are in bold. Xeric species are shaded.

| Species | Treatment | Average $T_{air}$<br>slope | 95% CI | Heatwave<br>$T_{air}$ slope | 95% CI |
| --- | --- | --- | --- | --- | --- |
| *Acronychia oblongifolia* | Control | 0.99 | 0.90-1.10 | **1.62** | **1.44-1.80** |
| *Pomaderris apetala* | Control | 0.96 | 0.86-1.05 | 0.98 | 0.85-1.12 |
| *Brachychiton rupestris* | Control | 1.00 | 0.96-1.04 | **1.14** | **1.10-1.19** |
| *Pittosporum angustifolium* | Control | 1.00 | 0.93-1.07 | **1.40** | **1.31-1.49** |
| *Acronychia oblongifolia* | Drought | 0.97 | 0.87-1.07 | **1.59** | **1.42-1.76** |
| *Pomaderris apetala* | Drought | 1.00 | 0.85-1.15 | **1.33** | **1.07-1.58** |
| *Brachychiton rupestris* | Drought | 1.00 | 0.85-1.15 | **1.47** | **1.29-1.66** |
| *Pittosporum angustifolium* | Drought | 1.00 | 0.92-1.08 | **1.53** | **1.41-1.65** |

Stomatal decoupling was clearly observed in droughted *P. angustifolium* plants (Supporting Information Fig. S6B), but evidence was weaker for droughted *A. oblongifolia* plants (Supporting Information Fig. S6A). The g_s_ of droughted plants did not change during the heatwave for either species (*P. angustifolium*: *F*_1,39_=0.57, *p*=0.458; *A. oblongifolia*: *F*_1,38_=1.72, *p*=0.198), but the associated decrease in A_net_ at high air temperature was only significant for *P. angustifolium* (*F*_1,39_=13.65, *p*<0.001; *A. oblongifolia*: *F*_1,38_=0.11, *p*<0.737; Supporting Information Fig. S5). Again, the observed g_s_ during heatwave conditions was greater than that predicted by the USO fitted to droughted plants under average summer temperatures (Supporting Information Fig. S6A, S6B) – the slope of the relationship between observed and predicted g_s_ was about 1.5 for both *P. angustifolium* and *A. oblongifolia* (Table 3).

There was little evidence for stomatal decoupling in the xeric species *Brachychiton rupestris*, and even less for the rainforest species *Pomaderris apetala*. High-temperature responses between A_net_ and g_s_ were coupled for both species – as expected from the decline in A_net_ at high air temperature (*P. apetala*: *F*_1,39_=36.11, *p*<0.001; *B. rupestris*: *F*_1,40_=11.11, *p*=0.002; Supporting Information Fig. S5), the g_s_ of control plants significantly decreased during the heatwave for *P. apetala* (*F*_1,39_=13.35, *p*<0.001; Fig. 4C) and *B. rupestris* (*F*_1,40_=8.25, *p*=0.007; Fig. 4D). Similarly, both A_net_ and g_s_ of droughted plants decreased during the heatwave for *P. apetala* (A_net_: *F*_1,33_=85.90, *p*<0.001; g_s_: *F*_1,33_=31.90, *p*<0.001; Supporting Information Fig. S5, S6C) and *B. rupestris* (A_net_: *F*_1,34_=53.06, *p*<0.001; g_s_: *F*_1,34_=22.49, *p*<0.001; Supporting Information Fig. S5, S6D). The observed g_s_ during heatwave conditions did not differ from that predicted by the USO fitted to control plants under average summer temperatures for *P. apetala*, with a slope of about 1 (Fig. 4C), although there was a small increase in slope to 1.14 for *B. rupestris* (Table 3, Fig. 4D). The observed g_s_ during heatwave conditions was higher than that predicted by the USO fitted to droughted plants under average summer temperatures – the slope of the relationship between observed and predicted g_s_ was 1.33 (Supporting Information Fig. S6C) for *P. apetala* and 1.47 for *B. rupestris* (Table 3, Supporting Information Fig. S6D).

### Temperature effect on minimum conductance

The temperature response of g_min_ depended on species (T_air_ × species: *F*_3,205_=24.43, *p*<0.001). There was a clear biphasic temperature response on *g*_min_ of xeric *P. angustifolium*, where there was an increase in slope at a breakpoint near 34 °C (*r*^2^=0.46, *p*<0.001; Fig. 5B). The two slopes between temperature and g_min_ were 0.002 (−0.022 to 0.027, 95% confidence interval) and 0.095 (0.016-0.175, 95% confidence interval). There was no relationship between temperature and g_min_ for *A. oblongifolia*, *P. apetala*, or *B. rupestris* (Fig. 5). Species differences in mean g_min_ existed (*F*_3,205_=66.96, *p*<0.001) – the xeric *B. rupestris* had the lowest g_min_ (0.001 mol m^-2^ s^-1^), and the mesic *P. apetala* had the highest g_min_ (0.004 mol m^-2^ s^-1^), when compared at T_air_ <35 °C (Fig. 5). The other two species had similar intermediate mean g_min_ (0.003 mol m^-2^ s^-1^; Fig. 5).

**Figure 5.**
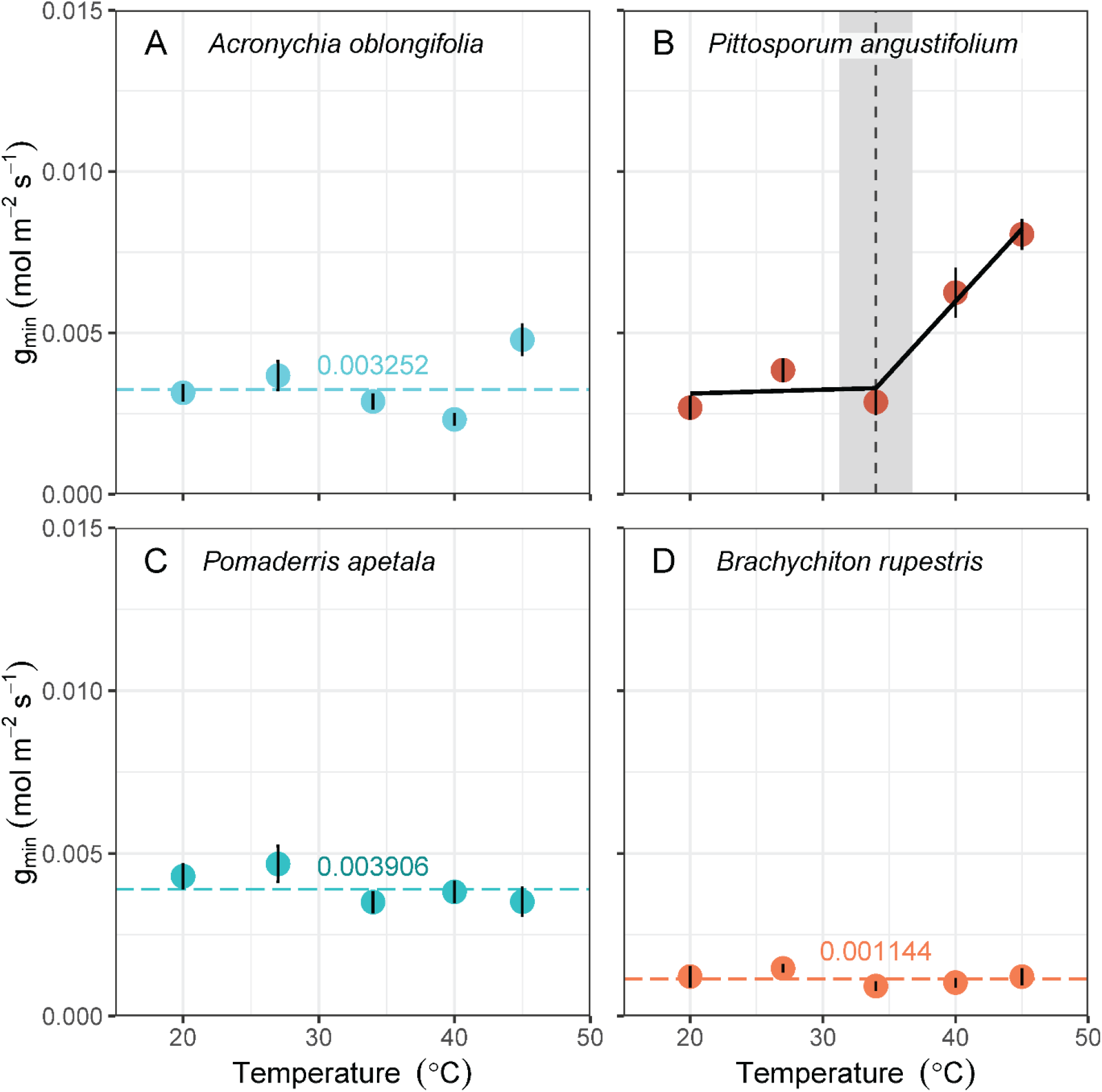
The effect of temperature on minimum conductance (g_min_, mol m^-2^ s^-1^) among four tree species from rainforests (cyan) or semi-arid woodlands (coral). Values are means (±SE) of 5‒15 leaves per temperature. Horizontal dashed lines indicate species mean if temperature had no effect on g_min_. The solid black line illustrates the significant model of best fit (piece-wise regression, as determined using Akaike’s Information Criterion, corrected for small sizes; AICc). Vertical dashed line and grey bars indicate statistically significant breakpoints and their 95% confidence intervals, respectively.

### Temperature effect on stomatal kinetics

Temperature significantly affected many of the stomatal kinetics parameters associated with light-induced stomatal opening, including g_low_ (*F*_1,46_=8.60, *p*=0.007), g_op_ (*F*_1,46_=12.80, *p*=0.001), and the slope parameter [Δg_s_/Δt]_max_ (*F*_1,44_=12.94, *p*=0.001) but not t_50_ (*F*_1,46_=0.05, *p*=0.819). Temperature significantly affected the dark-induced stomatal closing parameter t_50_ (*F*_1,39_=36.11, *p*<0.001) but not the slope parameter [Δg_s_/Δt]_max_ (*F*_1,39_=0.63, *p*=0.437). When averaged across species, maximum rates of stomatal opening were higher at 34 °C than at 25 °C (*t*_59_=3.89, *p*<0.001), while stomata closed at similar lower rates regardless of temperature (*t*_59_=0.84, *p*=0.402; Fig. 6A). The t_50_, or time taken to reach halfway between g_low_ and g_op_, was longer for stomatal closing at 34 °C than at 25 °C (*t*_59_=6.77, *p*<0.001), requiring 30 and 16 minutes, respectively, but the t_50_ of stomatal opening was similar at both temperatures, when averaged across species (*t*_61_=0.61, *p*=0.542; Fig. 6B).

**Figure 6.**
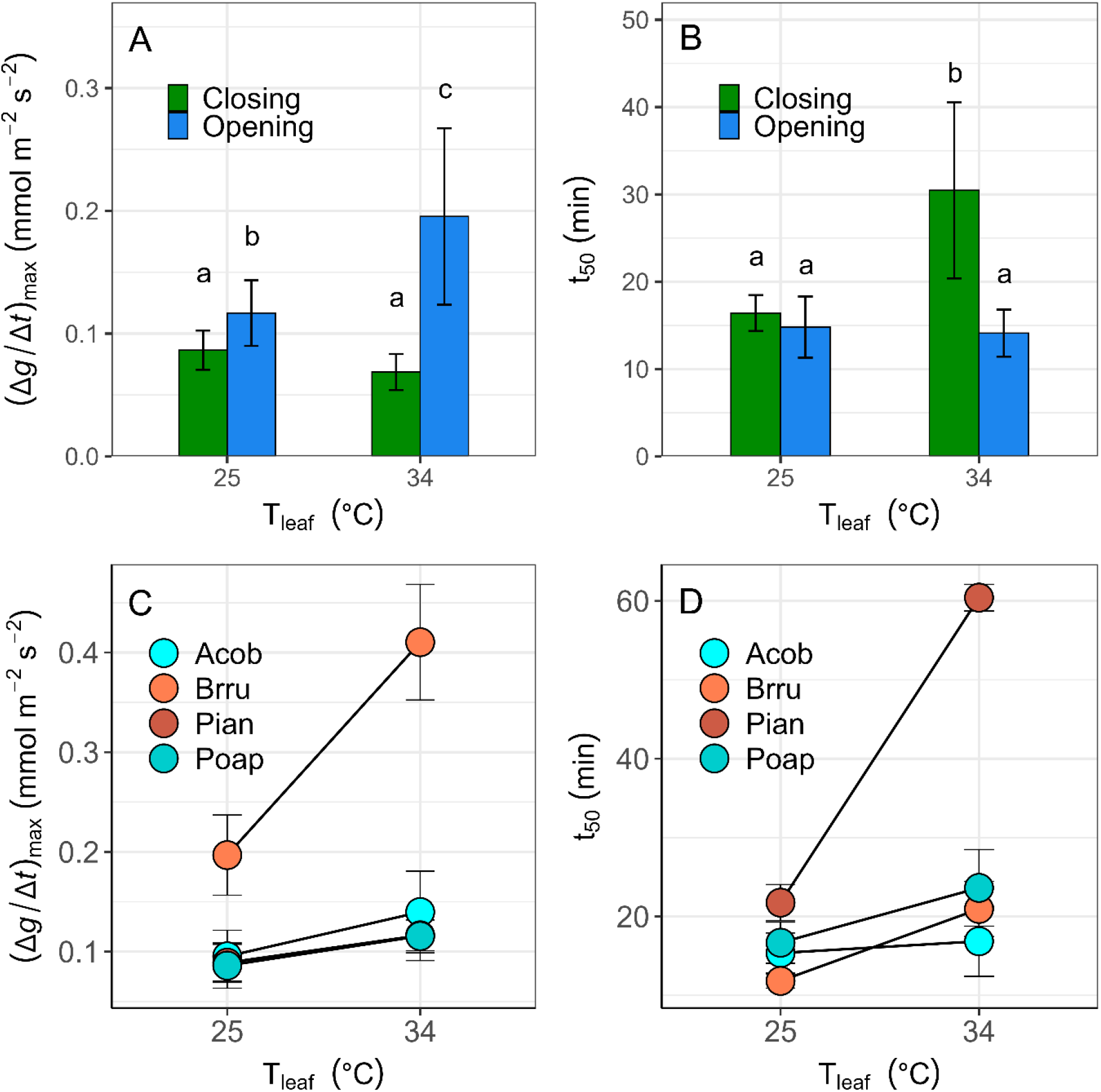
The effect of temperature on stomatal kinetics parameters including the (A) maximum rate of change in g_s_ ([Δg_s_/Δt]_max_) and (B) time taken to reach 50% of the point between g_low_ and g_op_ (t_50_) across all four species. Stomatal opening is light induced, while stomatal closing is dark induced. Means not connected by the same letter are significantly different (Tukey honestly significant difference, *p*<0.05). The main effect of temperature was significant, but temperature responses depended on species (Acob: *Acronychia oblongifolia*, Pian: *Pittosporum angustifolium*, Poap: *Pomaderris apetala*, Brru: *Brachychiton rupestris*), as shown for (C) [Δg_s_/Δt]_max_ during light-induced stomatal opening and (D) t_50_ during dark-induced stomatal closing.

The temperature response of g_low_, g_op_, and [Δg_s_/Δt]_max_ depended on species (T_leaf_ × species: *F*_3,46_≥3.12, *p*≤0.044). The stomatal opening kinetics of the most isohydric species *B. rupestris* were more sensitive to temperature, relative to the other species (Fig. 6C). Leaves of *B. rupestris* at a typical summer T_leaf_ of 34 °C had 2.5 times higher g_low_, 1.7 times higher g_op_, and over two times the [Δg_s_/Δt]_max_, relative to the standard T_leaf_ (25 °C; Fig. 6C). For other species, temperature-induced increases were more modest for all stomatal opening parameters, ranging from 1.2–1.4 times higher (Fig. 6C).

The temperature response of t_50_ for dark-induced stomatal closing also depended on species (T_leaf_ × species: *F*_3,39_=14.06, *p*<0.001). The most anisohydric species *P. angustifolium* had the slowest stomata to open (*F*_3,46_=6.17, *p*=0.005) and close (*F*_3,39_=21.13, *p*<0.001) – its t_50_ for opening was 24 ± 5 min, significantly higher than 10 ± 0.5 min in *A. oblongifolia* and *B. rupestris*. Stomatal closing of *P. angustifolium* took 2.6 times longer at 34 °C than at 25 °C, while other species had t_50_ that was about 1.1–1.8 times longer at 34 °C than at 25 °C (Fig. 6D).

## Discussion

Evidence of stomatal decoupling at high T_leaf_ for forest trees is mounting (Drake *et al*., 2018; Aparecido *et al*., 2020; Marchin *et al*., 2023; Gauthey *et al*., 2024), but the response is highly variable across species (Wang *et al*., 2026). We found stomatal decoupling at high T_leaf_ occurred in two of four tree species, including one rainforest (*A. oblongifolia*) and one xeric species (*P. angustifolium*), albeit mean A_net_ remained above zero for most study plants (Fig. 4, Table 3, Supporting Information Fig. S5, S6). One limitation of our study is that the temperature response of g_s_ and g_min_ are confounded with VPD, as our experimental heatwave caused an increase in both temperature and VPD, albeit daytime relative humidity was only marginally higher by 4% during the heatwave (Supporting Information Fig. S1). Hereafter, we refer to ‘the temperature response’ for simplicity but, in fact, this is a combined temperature and VPD response. Instead of disentangling the temperature and VPD effects on g_s_ and g_min_, we focussed on species differences in plant thermal and hydraulic balance and plausible biophysical mechanisms for stomatal decoupling.

### Plant water use, thermal regulation, and vulnerability to heatwaves across species

Both xeric species (*B. rupestris*, *P. angustifolium*) were tolerant to experimental stress, preventing their leaves from overheating (Fig. 2) and avoiding crown dieback (Fig. 3), although each used different strategies of water use and thermal regulation. Seedlings of the Australian bottletree *B. rupestris* had many drought-avoiding traits, matching previous findings (Reynolds *et al*., 2018) – the highest A_sat_ (Table 2), lowest wood density (Table 1), isohydric behaviour of closing stomata under mild drought stress (Fig. 1A), and slow recovery of g_s_ after compound heat and drought stress (Supporting Information Fig. S4). It also had the fastest stomatal kinetics with the highest temperature sensitivity for light-induced stomatal opening (Fig. 6C), and little evidence of stomatal decoupling (Table 3, Fig. 4, Supporting Information Fig. S5, S6) along with the lowest *g*_min_ (0.001 mol m^-2^ s^-1^), which was not sensitive to increasing temperature (Fig. 5D). As a result, *B. rupestris* maintained the highest Ψ_mid_ (Fig. 1C) and the largest thermal safety margins (i.e., T_leaf_ << T_crit_) under drought and heat stress (Fig. 2D), with no leaf shedding observed. These traits suggest *B. rupestris* has an opportunistic leaf gas exchange strategy that maximises carbon gain during rare, unpredictable rainfall events (Reynolds *et al*., 2018), while effectively buffering against water stress through tight isohydric stomatal behaviour, little stomatal decoupling, fast stomatal kinetics, an impermeable leaf cuticle, and if those safeguards fail, leaf shedding to limit water loss.

The xeric species *P. angustifolium* was the most anisohydric species studied here, having the lowest π_tlp_ (Table 2) and allowing its Ψ_mid_ to surpass π_tlp_ under heat and/or drought stress (Fig. 1A). In contrast to *B. rupestris*, prevention of overheating in *P. angustifolium* was strongly tied to its high capacity for transpirational cooling and leaf thermoregulation (Table 2, Fig. 1) – both g_s_ and g_min_ were sensitive to temperature, along with the strongest decoupling response (Table 3, Fig. 4, Supporting Information Fig. S5, S6) and g_min_ above 5 mmol m^-2^ s^-1^ when T_leaf_ exceeded 35 °C (Fig. 5B). Similar findings of leaf cooling via transpiration have been reported to promote survival of riparian cottonwood (*Populus fremontii*) saplings in the desert (Posch *et al*., 2024). Transpiration likely helped to facilitate photosynthesis in *P. angustifolium* (Supporting Information Fig. S5), aided by heat-tolerant leaf physical traits (e.g., high LMA, narrow leaves; Table 1) that buffered leaves against thermal stress (Leigh *et al*., 2012). Surprisingly, *P. angustifolium* had the slowest stomatal kinetics of all species in terms of half-times (t_50_ ≥20 min; Fig. 6D), although its maximum rate of change in g_s_ ([Δg_s_/Δt]_max_) was similar to the two rainforest species (Fig. 6C). In summary, our results suggest leaf thermoregulation in *P. angustifolium* at high T_leaf_ resulted from a coordinated temperature response involving g_s_, g_min_, and stomatal kinetics.

The two rainforest species were both more vulnerable to cumulative stress, particularly *P. apetala*, whose leaves surpassed their T_crit_ (Fig. 2C), resulting in moderate crown dieback (>20%) for three seedlings (Fig. 3A). Although *P. apetala* had the lowest T_crit_ under baseline conditions (Table 2), plants thermally acclimated to heat stress, achieving similar T_crit_ to other species by the fifth heatwave day. This acclimation was insufficient to avoid damage, but notably, stomatal decoupling was not observed in *P. apetala* (Table 3, Fig. 4, Supporting Information Fig. S5, S6). These results suggest stomatal decoupling may not occur in all taxa (Aparecido *et al*., 2020; Marchin *et al*., 2023) but could be restricted to species with high capacity for transpirational cooling (5-6 °C; *A. oblongifolia*, *P. angustifolium*), given the two species with low transpirational cooling capacity (∼2 °C; *P. apetala*, *B. rupestris*) had little decoupling (Table 2). Species with high transpiration capacity have been previously shown to have higher vein densities and stomatal pore areas, whereas stomatal size and density were not consistently related with transpiration capacity (Lin *et al*., 2017). Although there is logic to the hypothesis that stomatal decoupling is restricted to species with high transpiration capacity, another factor may have caused the differential decoupling response across species – stomatal decoupling may require full sunlight (i.e., PAR >1000 µmol m^-2^ s^-1^).

Despite attempting to standardise conditions during our experiment, both species with stomatal decoupling (*A. oblongifolia*, *P. angustifolium*) were exposed to greater PAR during their experimental heatwave (Supporting Information Fig. S3), resulting in heat-induced chronic photoinhibition (negative ΔF_v_/F_m_; Fig. 3B). There was no evidence for photoinhibition in the species without stomatal decoupling (*P. apetala*, *B. rupestris*; Fig. 3B), likely due to cloudy conditions and lower PAR (<500 µmol m^-2^ s^-1^; Supporting Information Fig. S3) during the last three days of the experimental heatwave (i.e., when F_v_/F_m_ was measured). A process-based model of g_s_ (Buckley *et al*., 2003) predicted weak stomatal responses to temperature under low light but strong temperature responses under high light, when VPD was held constant to isolate the influence of photosynthetic processes on g_s_ (Mills *et al*., 2024). Stomata may sense and respond to some measure of the balance between the Calvin cycle and the light reactions of photosynthesis, such as the redox state of plastoquinone in the electron transport chain (Busch, 2014), the ATP concentration in mesophyll cells (Farquhar & Wong, 1984; Buckley *et al*., 2003), or a vapour-phase signal that travels from mesophyll to guard cells (Sibbernsen & Mott, 2010). Our results suggest that stomatal decoupling may require high irradiance to trigger photoinhibition, but further research is needed.

### What drives stomatal decoupling at high temperature – g_s_ or g_min_?

The strong positive correlation between net photosynthesis (A_net_) and g_s_ (Wong *et al*., 1979) represents the fundamental plant trade-off between maximizing carbon gain and minimizing water loss (Cowan & Farquhar, 1977). This water use efficiency hypothesis forms the basis of a prominent group of g_s_ optimality models (Medlyn *et al*., 2011) that predicts leaf gas exchange in many land surface models, but it poorly captures leaf gas exchange at high T_leaf_ (Marchin *et al*., 2023; Sicangco *et al*., 2026), particularly when stomatal decoupling has occurred (e.g., *A. oblongifolia*, *P. angustifolium*; Table 3). More recent g_s_ models use an instantaneous optimisation approach, where stomatal behaviour is optimised to instantaneously maximise the difference between carbon gain and hydraulic cost (Wolf *et al*., 2016; Sperry *et al*., 2017). The key difference is that the water cost is allowed to vary over short time scales (i.e., days to weeks; (Wolf *et al*., 2016), which improves forest responses to drought (Sabot *et al*., 2020) and better predicts stomatal decoupling during heatwaves (Sicangco *et al*., 2026).

Previous studies that identified stomatal decoupling could not resolve whether accelerated leaf water loss occurred through open or closed stomata (Marchin *et al*., 2023; Gauthey *et al*., 2024; Gauthey *et al*., 2026). Here, we measured g_s_ and g_min_ on the same plants, finding that accelerated water loss at high T_leaf_ occurred through stomata for both species with decoupling, as g_s_ was an order of magnitude above maximum mean g_min_ (i.e., at 45 °C; Fig. 4, Supporting Information Fig. S6). Evergreen trees with long-lived leaves typically have low g_min_, with most species having values <5 mmol m^-2^ s^-1^ (Trueba *et al*., 2026), in agreement with our observations. In three of our study species, g_min_ did not increase at high temperature (Fig. 5A), but we did not control VPD, which has been shown to counteract and dampen the g_min_ temperature response (Garen *et al*., 2026). The lack of a temperature response of g_min_ in some species has been attributed to higher cuticular wax coverage, along with differences in wax chemistry that function to increase its melting point (Bueno *et al*., 2019). Our results suggest stomatal decoupling occurred primarily through open stomata (i.e., g_s_; Fig. 4, Supporting Information Fig. S6) but also by a smaller fraction of water lost through leaky stomata and the leaf cuticle, or g_min_, in xeric *P. angustifolium* (Fig. 5). Existing g_s_ models should be modified to more accurately represent the effects of high temperatures and VPD on plant-atmosphere exchange of carbon and water.

### How does warming affect stomatal kinetics?

The stomata of intact leaves respond to light and CO_2_, partly mediated by temperature-induced changes in leaf photosynthetic processes (Lawson *et al*., 2018). The light response of stomata is mediated by photoreceptor systems in the guard cell chloroplasts, involving induction of membrane ion transport and osmotic pressure changes that lead to turgor-driven changes in stomatal aperture (Zeiger, 1983). The biophysical mechanisms underlying stomatal responses to temperature remain unknown (Mills *et al*., 2024), along with if and how responses vary across species. To our knowledge, our study is the first to report that warming increased the maximum rate of change in g_s_ during light-induced stomatal opening ([Δg_s_/Δt]_max_; Fig. 6A) and lengthened the time required for dark-induced stomatal closing (Fig. 6B). Notably, stomatal response rates to light and VPD were similar in a C_3_ (soybean) and a C_4_ (sugarcane) plant, suggesting there may be a common mechanism controlling stomatal movements to both stimuli (Grantz & Zeiger, 1986). Stomatal opening is promoted by blue light, which is sensed by phototropin photoreceptors (Kinoshita *et al*., 2001) and activates plasma membrane H^+^-ATPases (Shimazaki *et al*., 1986) that cause hyperpolarisation and activation of potassium channels in guard cell membranes (Ilan *et al*., 1995). High temperature-induced stomatal opening in *Arabidopsis thaliana* involved phototropin photoreceptors, plasma membrane H^+^-ATPases, and 14-3-3 proteins, although this pathway has not been fully elucidated (Kostaki *et al*., 2020). Other biophysical mechanisms outside the guard cells are likely to be involved, such as modulation of leaf water status by temperature-dependent changes in water transport, abscisic acid metabolism, or temperature-driven shifts in photosynthetic processes (Mills *et al*., 2024), as described above.

The direction of stomatal kinetics temperature responses were consistent across species from diverging biomes, but the magnitude of responses varied substantially among species (Fig. 6). Stomatal kinetics of xeric species were more sensitive to temperature than rainforest species. For xeric *B. rupestris*, the maximum rate of light-induced stomatal opening was two times faster in leaves at 34 °C, relative to 25 °C (Fig. 6C). For xeric *P. angustifolium*, the half-time for dark-induced stomatal closure was 60 ± 2 minutes at 34 °C but only 22 ± 2 minutes at 25 °C (Fig. 6B). Our results suggest accelerated water loss at high T_leaf_ involves faster stomatal opening or slower stomatal closing in heat-tolerant species. Faster stomatal kinetics (i.e., high [Δg_s_/Δt]_max_) have previously been associated with smaller and more numerous stomata, at least among closely related tree species (Drake *et al*., 2013), which may offset the risk for larger water potential gradients associated with high gas exchange rates in environments with fluctuating light and evaporative demand (Drake *et al*., 2013; Meinzer *et al*., 2017). Our findings of slower stomatal closing with warming, along with the long stomatal response times found in xeric *P. angustifolium*, is suggestive of anisohydric species prioritising carbon gain over water use efficiency when water is available (Marchin *et al*., 2016; Wolf *et al*., 2016; Meinzer *et al*., 2017). Further research should test a wider range of temperatures to determine if stomatal kinetics respond linearly or non-linearly to increasing temperature, along with a wider range of species to determine if stomatal kinetics varies predictably across taxa.

## Conclusions

Stomatal decoupling at high T_leaf_ caused accelerated water loss in two of four species, suggesting that leaf cooling is a critical driver of stomatal regulation in the heat-stressed leaves of some species. These two species (*Acronychia oblongifolia*, *Pittosporum angustifolium*) had leaves with high capacity for transpirational cooling of 5-6 °C under summer temperatures (T_air_= 35 °C), whereas decoupling was not observed in two species (*Pomaderris apetala*, *Brachychiton rupestris*) with low capacity for transpirational cooling (∼2 °C) but also no photoinhibition, likely due to low photosynthetically active radiation during experimental imposition of peak heat stress. Leaf thermoregulation under heat stress occurred in one xeric species (*P. angustifolium*) but not the other (*B. rupestris*), and both anisohydric and isohydric strategies avoided overheating during drought through coordination of leaf physical traits with plasticity of leaf gas exchange, spanning temperature-induced changes in g_s_, g_min_, and stomatal kinetics. In contrast, leaf overheating and moderate crown dieback (>20%) occurred in the rainforest species *P. apetala*, despite strong thermal acclimation and cloudy conditions during peak heat stress. This suggests that some species prioritise acclimation of photosynthetic heat tolerance over leaf thermoregulation during heat stress, or alternatively, that full sunlight is needed to induce the stomatal decoupling response. Our results demonstrate that stomatal decoupling at high T_leaf_ occurred predominantly through open stomata (i.e., g_s_) rather than the leaf cuticle (i.e., g_min_), involving faster light-induced stomatal opening and slower dark-induced stomatal closing, especially in heat-tolerant species. Stomatal kinetics are an overlooked but informative trait that may help to better predict accelerated water loss under high T_leaf_ across species.

## Acknowledgements

This research was a contribution of the Which Plant Where project, which is funded by the Green Cities Fund, as part of the Hort Frontiers Strategic Partnership Initiative developed by Hort Innovation, with co-investment from Macquarie University, Western Sydney University, and the NSW Department of Planning and Environment, and funds from the Australian Government. RMM is the recipient of an Australian Research Council Discovery Early Career Researcher Award (project DE200100649) funded by the Australian Government; the views expressed herein are those of the authors and are not necessarily those of the Australian Government or the Australian Research Council. We thank Andrew Gherlenda of Western Sydney University for his technical assistance with the glasshouse facility and plant growth. We thank Nikki Bown, Agnieszka Wujeska-Klause, John Foster, and Samiya Tabassum for help in data collection and Leigh Staas for project administrative support.

## Author contributions

RMM: Conceptualisation, Investigation, Formal analysis, Writing – Original Draft; AO: Supervision, Writing – Review & Editing; MGT: Funding acquisition, Supervision, Writing – Review & Editing; DSE: Funding acquisition, Supervision, Writing – Review & Editing.

## Data availability

The data that support the findings of this study are available on request from the corresponding author.

