## Supporting Information for "Towards understanding stomatal decoupling under extreme heat: integrating stomatal kinetics, leaf light responses, and the role of stomatal versus cuticular conductance"

Renée M. Marchin[1], Alessandro Ossola[2,3], Michelle R. Leishman[3], David S. Ellsworth[1]

[1]Hawkesbury Institute for the Environment, Western Sydney University, Penrith, NSW, 2751, Australia

[2]Department of Plant Sciences, University of California Davis, Davis, California, USA

[3]Department of Biological Sciences, Macquarie University, North Ryde, NSW, 2109, Australia

The following Supporting Information is available for this article:

- Table S1
- Figures S1-S6

Table S1. Four native Australian tree species selected for this study, including their species code (used in figures), family, and climate of origin, which is described using the maximum temperature of the warmest month (°C) and precipitation of the driest month (mm). Climate data are temporal (from 1960–1990) and spatial (across natural occurrence range) means. Means not connected by the same letter are significantly different (Tukey honestly significant difference, $p<0.05$). Xeric species are shaded.

| **Species** | **Species code** | **Family** | **MTWM (°C)** | **PDM (mm)** |
|---|---|---|---|---|
| *Acronychia oblongifolia* (A.Cunn. ex Hook.) Endl. ex Heynh. | Acob | Rutaceae | 26.6 | 52 |
| *Pomaderris apetala* Labill. | Poap | Rhamnaceae | 20.5 | 54 |
| *Brachychiton rupestris* (T.Mitch. ex Lindl.) K.Schum. | Brru | Malvaceae | 32.2 | 24 |
| *Pittosporum angustifolium* Lodd. | Pian | Pittosporaceae | 31.3 | 17 |

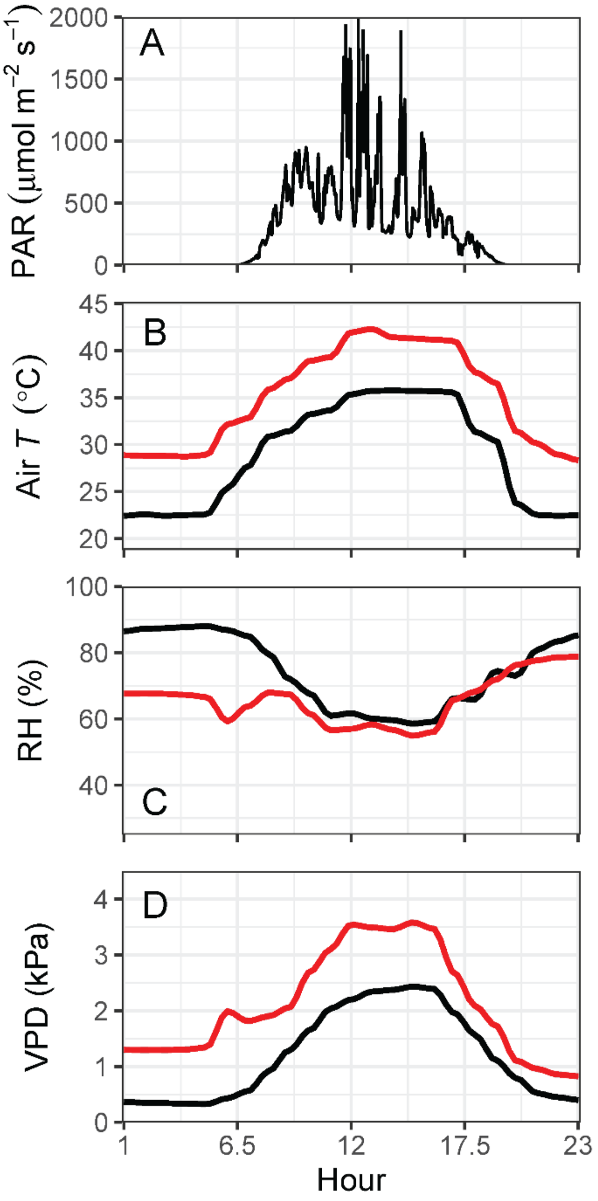


Figure S1. Environmental conditions inside the glasshouse, including (A) photosynthetically active radiation (PAR), (B) mean hourly air temperature, (C) mean hourly relative humidity, and (D) mean hourly vapor pressure deficit during baseline (black line) and heatwave conditions (red line). Data are shown for a representative sunny day (15 January 2019) in the middle of the experiment.

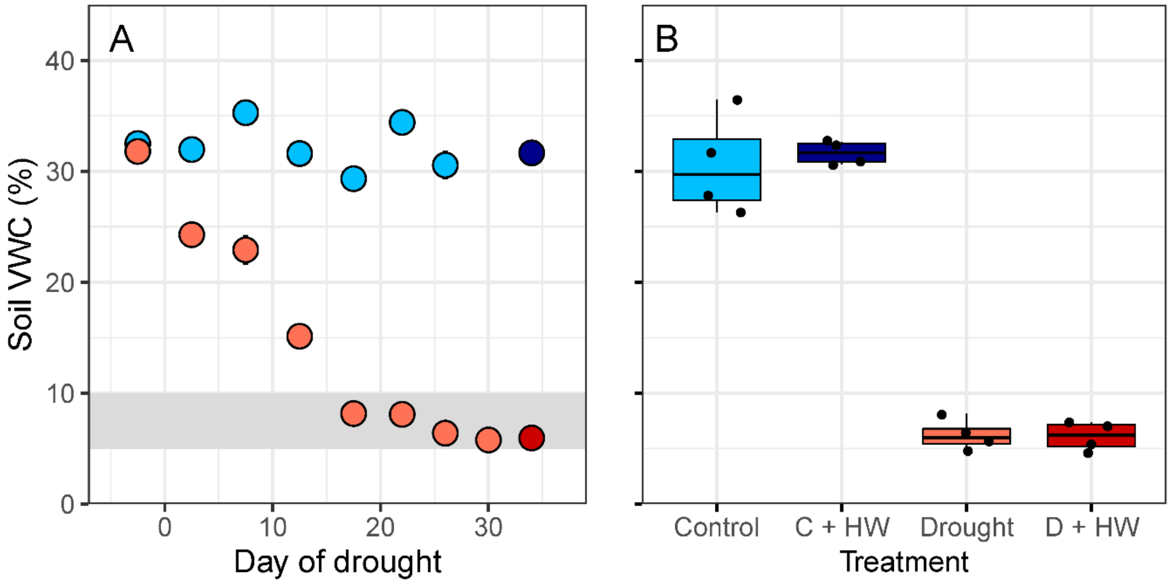


Figure S2. (A) Differences in mean soil volumetric water content (VWC, %) between control (blue) and drought (coral) pots over the five-week experimental drought period; error bars indicate SE but are smaller than the point size. The target drought intensity is shaded gray. Values are means of 20–40 plants per treatment; sample size varies with day due to binning of data collected on different days. (B) Mean soil VWC of four experimental treatments (Control, Drought, C + HW: Control + Heatwave, D + HW: Drought + Heatwave) measured during the fourth (baseline) and fifth (heatwave, HW) weeks of drought. Points are species means of 3–5 plants per treatment; Drought and D + HW are significantly lower than Control and C + HW treatments ($F_{3,71}$=413.17, $p$<0.001).

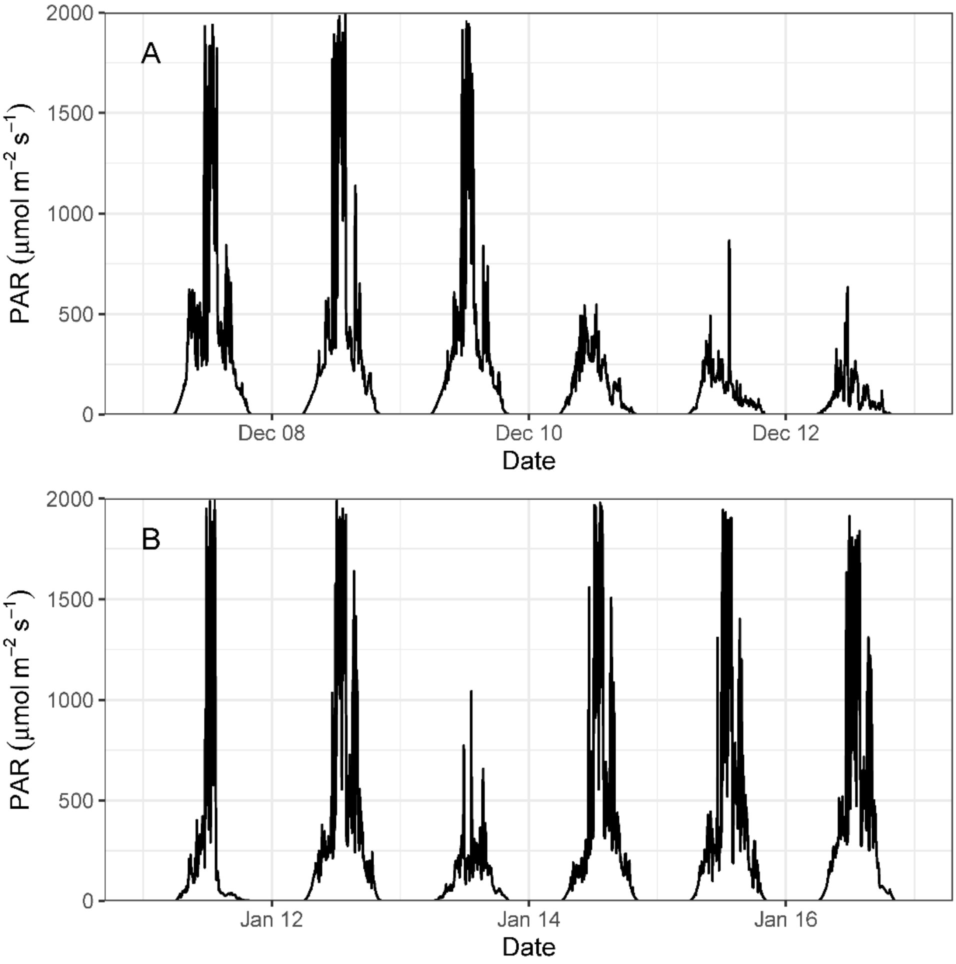


Figure S3. Photosynthetically active radiation (PAR) from (A) 7-12 December 2018, when *Brachychiton rupestris* and *Pomaderris apetala* were exposed to the experimental heatwave, and (B) 11-16 January 2019, when *Acronychia oblongifolia* and *Pittosporum angustifolium* were exposed to the experimental heatwave. Leaf gas exchange measurements during the heatwave were made on heatwave days 1, 3, and 5.

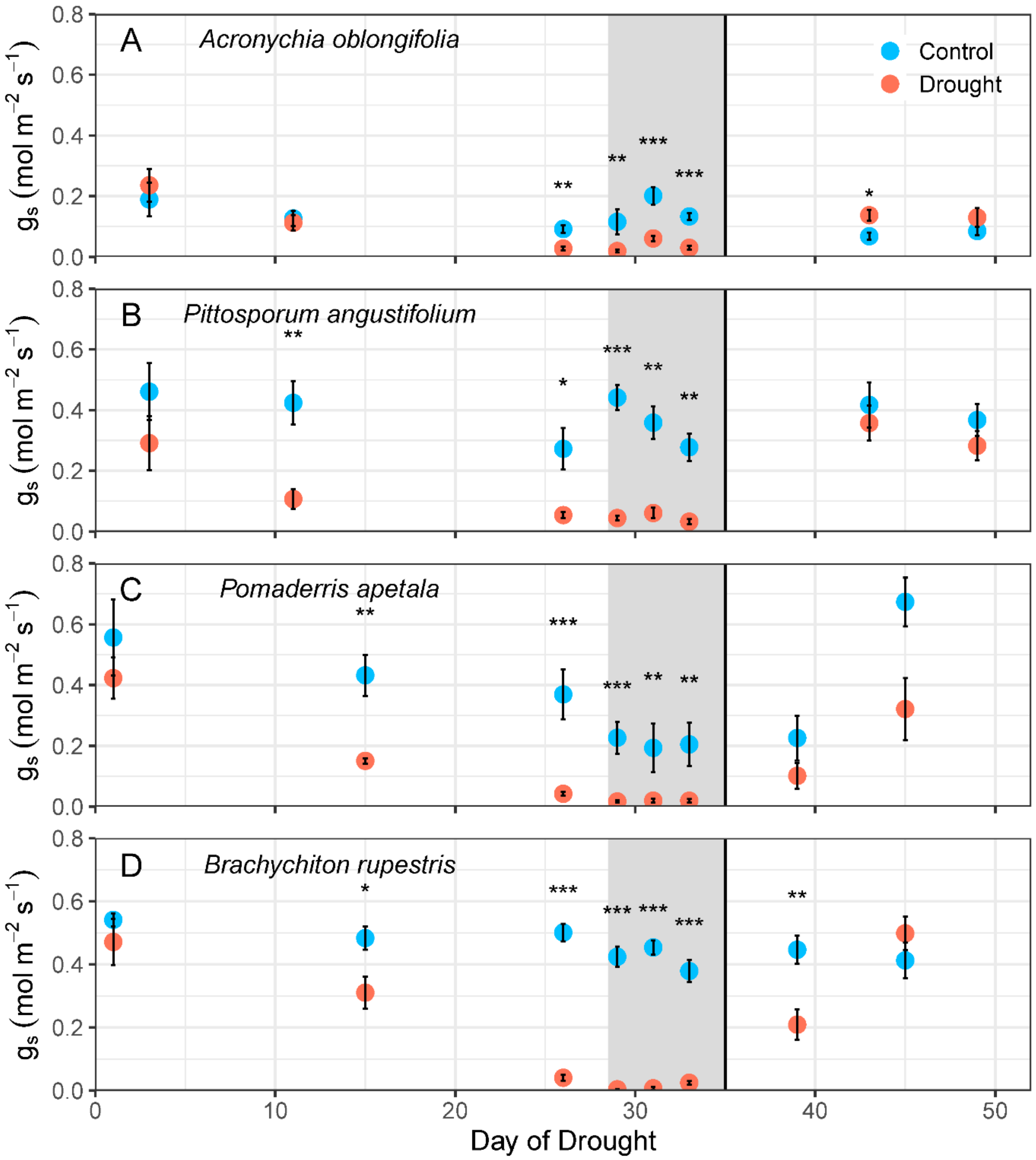


Figure S4. The effect of experimental drought on stomatal conductance ($g_s$) of (A) *Acronychia oblongifolia*, (B) *Pittosporum angustifolium*, (C) *Pomaderris apetala*, and (D) *Brachychiton rupestris*. Shaded region indicates when plants were exposed to the experimental heatwave; vertical line indicates when droughted plants were rewatered and allowed to recover. The four dates spanning drought days 26-33 were used to test for a heatwave effect on $g_s$ for all treatments. Points are means of five plants per species, where $g_s$ was averaged over 30-sec for two leaves of each plant; error bars indicate SE. Asterisks denote significant relationships: * $p$<0.05; ** $p$<0.01; *** $p$<0.001.

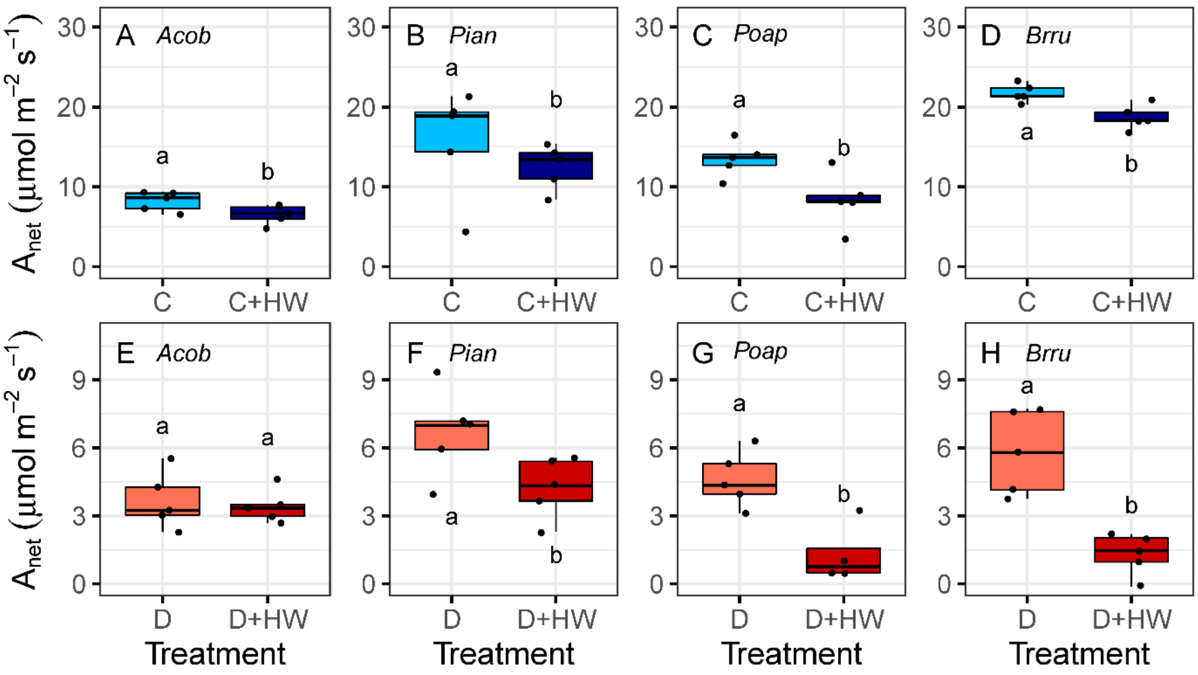


Figure S5. The effect of an experimental heatwave on mean leaf net $CO_2$ assimilation rate at light-saturation ($A_{net}$) for control and droughted plants of four tree species (Acob: *Acronychia oblongifolia*; Poap: *Pomaderris apetala*; Brru: *Brachychiton rupestris*; Pian: *Pittosporum angustifolium*). Points are means of two leaves per plant ($n$= 5 plants per treatment). Baseline data (C: Control, D: Drought) were measured two days before heatwave data; heatwave data (C + HW: Control + Heatwave, D + HW: Drought + Heatwave) includes measurements on heatwave days 1, 3, and 5. Means not connected by the same letter are significantly different (Tukey honestly significant difference, $p$<0.05).

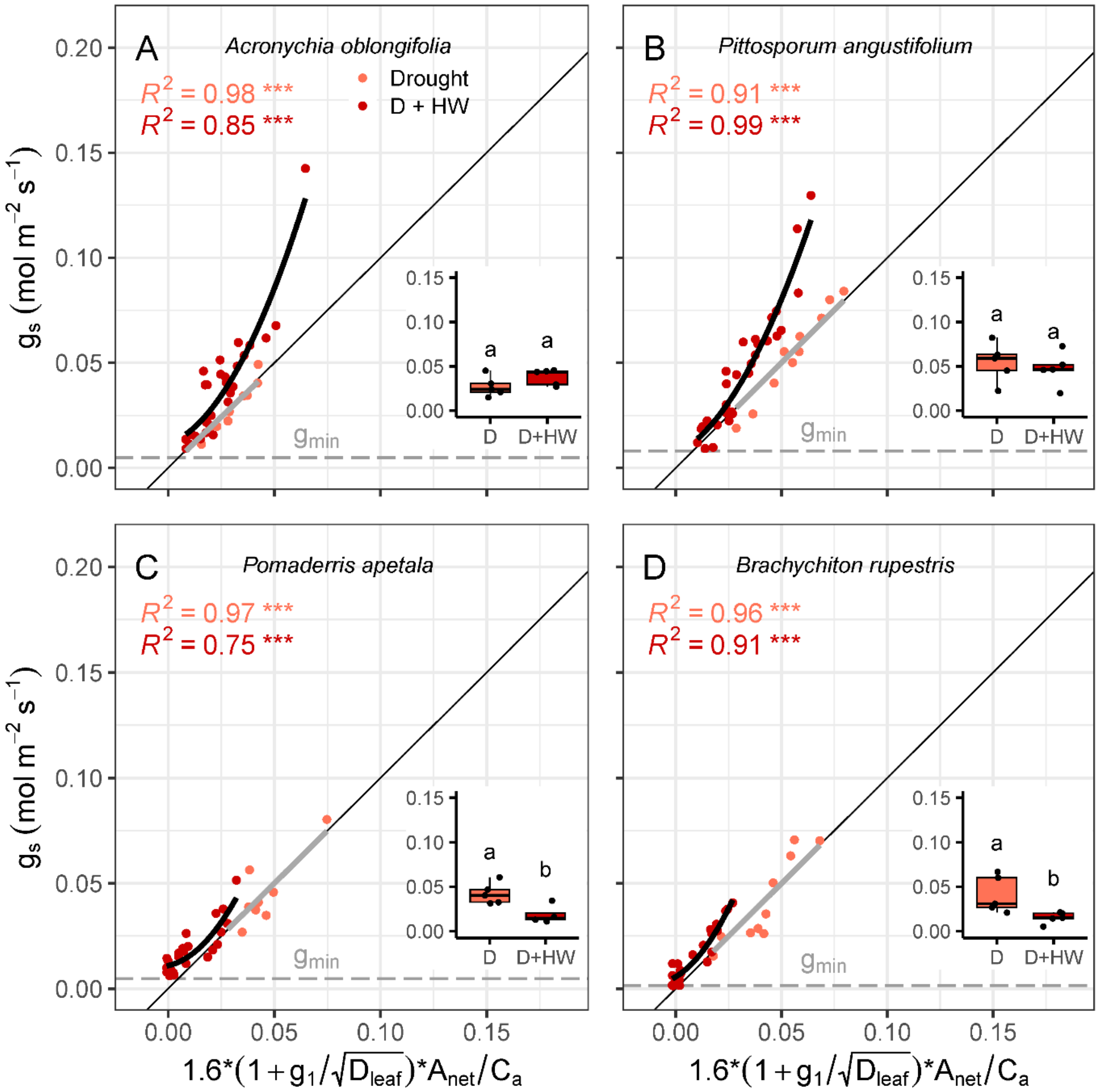


Figure S6. Relationships between observed stomatal conductance ($g_s$) and the parameterised unified stomatal optimisation model (USO) for drought plants of (A) *Acronychia oblongifolia*, (B) *Pittosporum angustifolium*, (C) *Pomaderris apetala*, and (D) *Brachychiton rupestris*. Responses are shown before (coral) and during the heatwave (red). Solid lines show linear or nonlinear regressions fit to the measured data. The 1:1 line between predicted and observed data is shown for reference. Points are 30-sec means for each leaf. Asterisks denote significant relationships: ** $p$<0.01; *** $p$<0.001. Insets show heatwave effects, where points are plant means. Means not connected by the same letter are significantly different (Tukey honestly significant difference, $p$<0.05). Maximum mean minimum conductance ($g_{min}$; horizontal dashed line) is shown for each species (i.e., at 45 °C for panels A and B, at 27 °C for panels C and D).
